# Wnt11/Fzd7 signaling compartmentalizes AKAP2/PKA to regulate L-type Ca^2+^ channel

**DOI:** 10.1101/741637

**Authors:** Kitti D. Csályi, Tareck Rharass, Maike Schulz, Mai H.Q. Phan, Paulina Wakula, Ketaki N. Mhatre, David Plotnick, Andreas A. Werdich, Henrik Zauber, Matthias D. Sury, Matthias Selbach, Frank R. Heinzel, Enno Klußmann, Daniela Panáková

## Abstract

Calcium influx through the voltage-gated L-type calcium channels (LTCC) mediates a wide range of physiological processes from contraction to secretion. Despite extensive research on regulation of LTCC conductance by PKA phosphorylation in response to β-adrenergic stimulation, the science remains incomplete. Here, we show that Wnt11, a non-canonical Wnt ligand, through its G protein-coupled receptor (GPCR) Fzd7 attenuates the LTCC conductance by preventing the proteolytic processing of its C terminus. This is mediated across species by protein kinase A (PKA), which is compartmentalized by A-kinase anchoring proteins (AKAP). Systematic analysis of all AKAP family members revealed AKAP2 anchoring of PKA is central to the Wnt11-dependent regulation of the channel. The identified Wnt11/AKAP2/PKA signalosome is required for heart development, controlling the intercellular electrical coupling in the developing zebrafish heart. Altogether, our data revealed Wnt11/Fzd7 signaling via AKAP2/PKA as a conserved alternative GPCR system regulating Ca^2+^ homeostasis.

---

The voltage-gated L-type calcium channel (LTCC) facilitates a major route for calcium entry into the cell. Consequently, its conductance must be strictly regulated, as calcium mediates a wide range of physiological processes, including those which can be electrically coupled, e.g. contraction, transcription, and secretion^1, 2^. Calcium influx via LTCC increases during membrane depolarization, and decreases upon calcium-and voltage-dependent inactivation^3, 4^. Regulation of LTCC conductance is most studied in the context of β-adrenergic receptor (β-AR)/PKA signaling^1, 5–7^.

The main pore-forming α1C subunit of the LTCC comprises cytosolic N-and C-terminal tails, and four transmembrane modules (Fig. 1a). Its long C-terminal tail (CT) serves as a scaffold for channel modulators, including kinases and phosphatases^3, 4^. Most of the α1C subunit exists in a truncated form^8^ due to calpain-dependent proteolytic processing of the CT^9, 10^. This results in a non-covalently associated distal CT, which inhibits the activity of the LTCC^10, 11^. This autoinhibition is alleviated upon stimulation of β-AR coupled to the stimulatory G protein, leading to cAMP-dependent phosphorylation of the CT by PKA^12^. PKA signaling is compartmentalized by AKAPs^13–15^, several of which have been identified to anchor PKA to the CT^16–21^. Whether PKA phosphorylation is absolutely required for LTCC regulation, however, has recently been brought into question^22, 23^. Moreover, which AKAPs are involved in a cell-and tissue-specific manner in this context remains inconclusive.

**Fig. 1.**
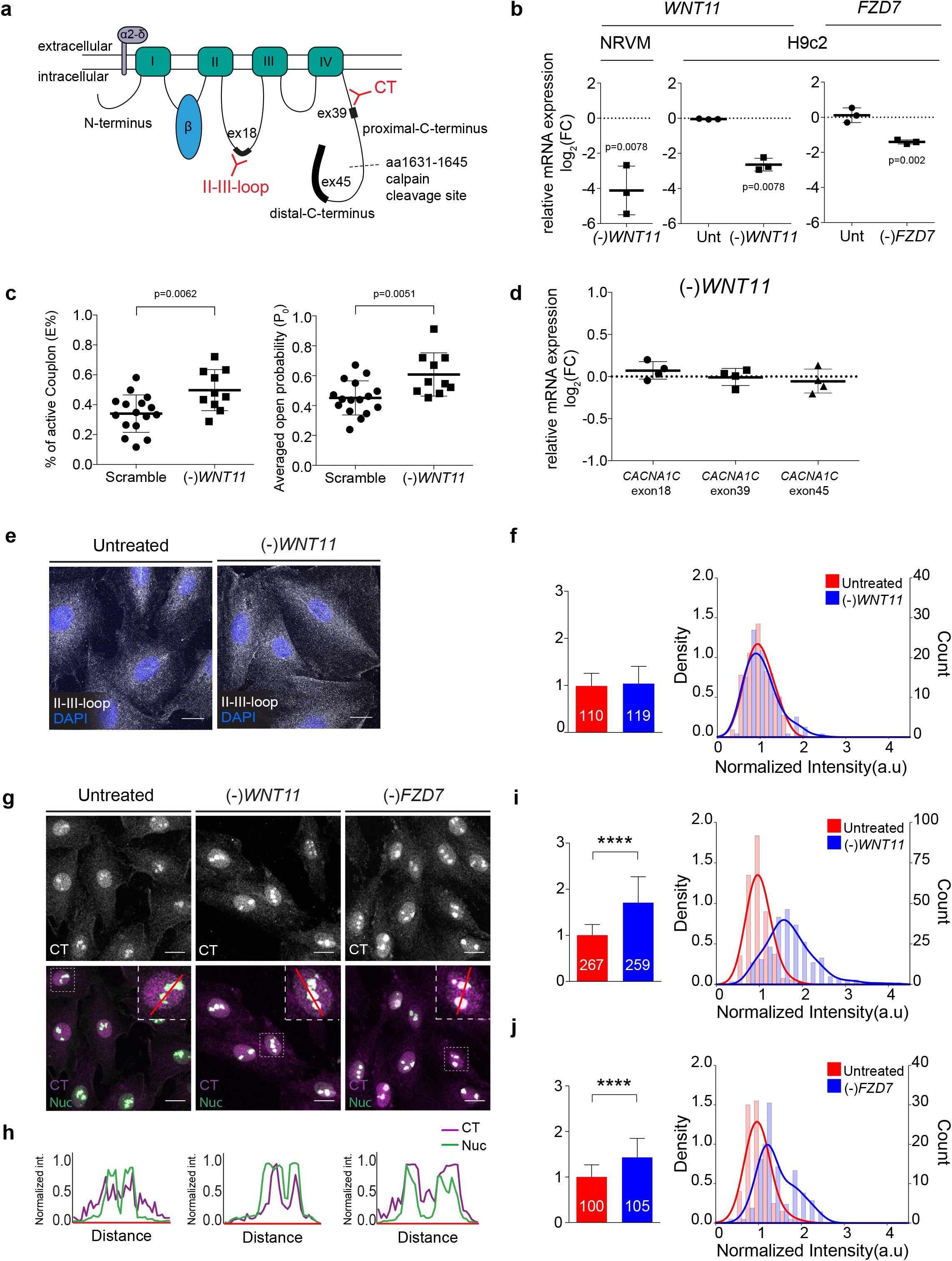
Wnt11/Fzd7 pathway regulates CT proteolysis. **a** Schematic of the L-Type Calcium channel. Calpain cleavage site (dotted black line), target sites for corresponding TaqMan probes (black), and antibody epitopes (red) are highlighted (reference sequence: *Ensembl* ID: ENSRNOT00000052017.6). **b** Quantification of relative mRNA expression of scramble and *WNT11* siRNA-treated NRVM (left), and untreated, scramble, *WNT11* and *FZD7* siRNA-treated H9c2 cells (right) showing siRNA efficacy. Data plotted as log_2_ of fold change (FC) ± SD of N = 3 experiments. Wilcoxon rank sum test, *WNT11*: *P*_Unt_>0.999, *FZD7*: *P*_Unt_*=*0.563. Unt: untreated, NRVM: neonatal rat ventricular myocytes. **c** Calcium release at LTCC-RyR couplons is higher in *WNT11* siRNA-transfected NRVM compared to scramble control. The percentage of line scanned with active couplons (left), and the average probability of calcium release at the couplons (right). Mean ± SD of N = 3 experiments, and n ≥ 10 cells. Unpaired t-test. **d** Quantification of relative mRNA expression of *CACNA1C* using Taqman probes targeting three different exons (as noted), in scramble and *WNT11* siRNA-transfected H9c2 cells. Data plotted as log_2_ of fold change ± SD of N = 3 experiments. Two-tailed Wilcoxon rank sum test, *P*_exon18_=0.13, *P*_exon39_ = 0.79, *P*_exon45_ = 0.233. **e** Representative maximum intensity Z-projection of untreated and *WNT11* siRNA-transfected H9c2 cells labeled with anti-II-III-loop (gray) and DAPI for nuclei (blue). **f** Quantified mean intensity of II-III-loop signals in whole cell. Bar graph (left) shows mean ± SD of N ≥ 3 experiments and n ≥ 110 cells as noted in the columns. Unpaired t-test with Welch’s correction, *P*=0.244. Corresponding histograms overlaid with density plots (right) show the distribution of normalized pixel intensity values within all analyzed images. **g** Representative maximum intensity Z-projection of CT signals in untreated, *WNT11* or *FZD7* siRNA-transfected H9c2 cells (gray scale, upper panel), and merged (magenta, lower panel) with nucleolin (Nuc) signals (green). Scale bar, 20 μm. **h** Line-scan analysis of CT and Nucleolin (Nuc) signals in ROIs in (G). (**i-j**) Quantified mean intensity of CT signals in nucleoli. Bar graphs (left) show mean ± SD of N ≥ 3 experiments and n ≥ 100 cells. Unpaired t-test with Welch’s correction, \*\*\*\**P*<0.0001. Corresponding histograms overlaid with density plots (right) show the distribution of normalized pixel intensity values within all analyzed images.

Wnt signaling is an evolutionarily conserved pathway within metazoans. Wnt ligands act through several receptors and co-receptors, most notably via the Fzd family of receptors belonging to Class F GPCRs^24, 25^. While structurally different from Class A receptors including β-AR ^24, 26^, the concept that Fzds do signal through heterotrimeric G proteins gained traction in recent years^25, 27–31^. The identification of the signaling components downstream of Fzd-G protein coupling, however, remains unresolved.

To date less attention has been paid to the regulation of LTCC via alternative GPCR systems. Wnt11 also regulates the LTCC conductance^32^, yet the underlying mechanism is unclear. In the present study, we show that Wnt11 signaling attenuates LTCC conductance by preventing proteolytic processing of the CT. Our data reveal that unlike the β-AR stimulation, the signal transduction initiated by Wnt11 ligand binding to Fzd7 is coupled to the inhibition of PKA. Furthermore, we identify the AKAP2/PKA signalosome playing a pivotal and conserved role in the Wnt11-dependent generation of the CT isoform. Tight regulation of AKAP2/PKA signalosome is required for Ca^2+^ homeostasis, heart morphogenesis, and controls the intercellular electrical coupling in the developing heart. AKAP/PKA compartmentalization by Wnt/Fzd G protein-coupled signaling may provide specificity towards different effectors of this complex conserved pathway.

## Results

### Wnt11 regulates CT proteolysis

To investigate the role of Wnt11 signaling in LTCC attenuation, we used a rat cardiomyoblast cell line (H9c2) and neonatal rat ventricular myocytes (NRVM). We confirmed expression of all the required components of the Wnt11 non-canonical pathway and the LTCC α1C subunit in H9c2 cells (Supplementary Fig. 1a). By siRNA treatment, we could efficiently reduce the expression levels of *WNT11* and *FZD7* in both cell models (Fig. 1b). Importantly, H9c2 cells and NRVMs responded to the loss of *WNT11* in the same manner as cardiomyocytes in the developing zebrafish heart^32^, including an increase in Connexin 43 (CX43) levels as shown by immunostaining and Western blotting (Supplementary Fig. 1b, c). Moreover, in *WNT11* siRNA-treated NRVM cells, the percentage (E%) of active LTCC-Ryanodine receptor (RyR) couplons, and their average open probability (P_0_), which directly relates to calcium release levels^33^, were significantly increased compared to scramble siRNA-treated cells (Fig. 1c). Collectively, these data suggested Wnt11 plays an essential role in regulating calcium homeostasis across species, and that our cell-based models are suitable to study the molecular mechanisms of Wnt11 in regulating LTCC conductance.

The transcription of the α1C subunit, encoded by the *CACNA1C* gene, is regulated through at least 4 cryptic promoters; transcription from three of these gives rise to functional proteins^34^. To determine whether Wnt11 signaling transcriptionally regulates the LTCC, we quantified relative mRNA expression of *CACNA1C* using 3 different TaqMan probes targeting exon 18, 39 and 45 covering all predicted transcripts (Fig. 1a), and corresponding to the II-III loop, proximal-CT (pCT), and distal-CT (dCT) of the α1C subunit (Fig. 1a). Expression levels of *CACNA1C* were unaltered in the absence of *WNT11* for all predicted transcripts (Fig. 1d).

To determine the LTCC localization within cardiomyoblasts, we first verified the efficiency of *CACNA1C* siRNA to test the specificity of the two different antibodies (Supplementary Fig. 1d, e): one recognizing the full-length α1C subunit (anti-II-III-loop), the other binding to the CT (anti-CT) (Fig. 1a). Lack of Wnt11 signaling had no effect on the localization or abundance of the α1C subunit (Fig. 1e, f). We found that the CT localized to the nucleus and the cytoplasm in both untreated and scramble siRNA-transfected cells (Fig. 1g). This behavior of the CT has been observed in myocytes, where its appearance in the nucleus is regulated developmentally, and by intracellular calcium^35^. Noteably, we also observed CT accumulating in a punctate pattern within the nucleus (Fig. 1g). Co-labeling with nucleolin (anti-Nuc) revealed CT specific nucleolar localization (Fig. 1g, h). The loss of *WNT11* induced CT nucleolar accumulation in both H9c2 (Fig. 1g-i) and NRVM cells (Supplementary Fig. 1f), as indicated not only by the increase in mean fluorescence intensities but also by a major shift in frequency distribution of fluorescence intensities. We focused on this particular phenomenon throughout this study.

Wnt11 together with its putative receptor Frizzled-7 (FZD7)^36, 37^ can signal through G_i_/G_0_^31, 38^ as well as through G_s_ proteins^27^. Importantly, the loss of *FZD7* also increased CT nucleolar localization (Fig. 1g, h, j). Taken together, our data indicated that Wnt11 regulates the proteolytic processing of the LTCC via its FZD7 receptor, prompting us to hypothesize that on the molecular level, Wnt11 might attenuate the LTCC through GPCR activity, akin to regulation by the β-AR/G_i_ protein-coupled system.

### Wnt11 pathway regulates CT formation via anchored PKA signaling

As PKA activation is the essential to cAMP signaling downstream of GPCR including β-AR, we tested whether Wnt11 may regulate PKA activity via GPCR-mediated signaling by utilizing the well-characterised PKA reporter AKAR4-NES. This genetically-encoded Förster resonance energy transfer (FRET) biosensor detects changes in cytosolic cAMP-dependent PKA activity as an increase in the yellow/cyan emission ratio^39^. *WNT11* siRNA-treated cells showed a significant increase in cytosolic PKA activity compared to untreated cells, as noted by an increase in normalized FRET emission (Fig. 2a). This result indicated Wnt11/Fzd7 signaling inhibits PKA activity, most likely through G_i_ coupling. Similar to dCT generation, which is well characterized upon β-AR/G_s_ stimulation^16, 21^, we first showed that the formation of the CT fragment depends on calpain proteolysis, which is independent of *WNT11* siRNA treatment (Supplementary Fig. 2a). We then asked whether CT formation, like dCT, could be induced by the β-AR/G_s_ system. Indeed, treatment with the β-AR agonist isoproterenol (ISO) effectively increased the CT generation (Fig. 2b, c). This suggested CT formation might be induced via PKA-dependent phosphorylation, similarly to dCT isoform formation. To study whether there is a crosstalk between Wnt11 and PKA signaling in regulating the CT isoform, we performed a series of pharmacological treatments. We reasoned that if Wnt11 does indeed regulate the LTCC via the PKA pathway, blocking PKA signaling should revert the loss of the Wnt11 effect. The direct activator of adenylyl cyclases, forskolin (FSK), elevated the level of phosphorylated PKA substrates, while treatment with the PKA inhibitor, Protein-kinase inhibitor (PKI), reduced phosphorylation of some substrates (e.g. 130kDa) (Supplementary Fig. 2b). FSK treatment strongly increased CT accumulation within the nucleoli (Fig. 2b, d), indicating that cAMP/PKA signaling is involved in the generation of this CT isoform. PKI treatment of *WNT11* siRNA-transfected cells reduced CT signals in the nucleoli, and shifted the frequency distribution of fluorescence intensities towards those of control cells (Fig. 2b, e). This was corroborated by the treatment of *WNT11* siRNA-treated cells with H89, a broader protein kinase inhibitor (Supplementary Fig. 2a), while neither okadaic acid, a phosphatase inhibitor, nor DMSO, the solvent of our pharmacological agents, reversed the loss of the Wnt11 effect (Supplementary Fig. 2a).

**Fig. 2.**
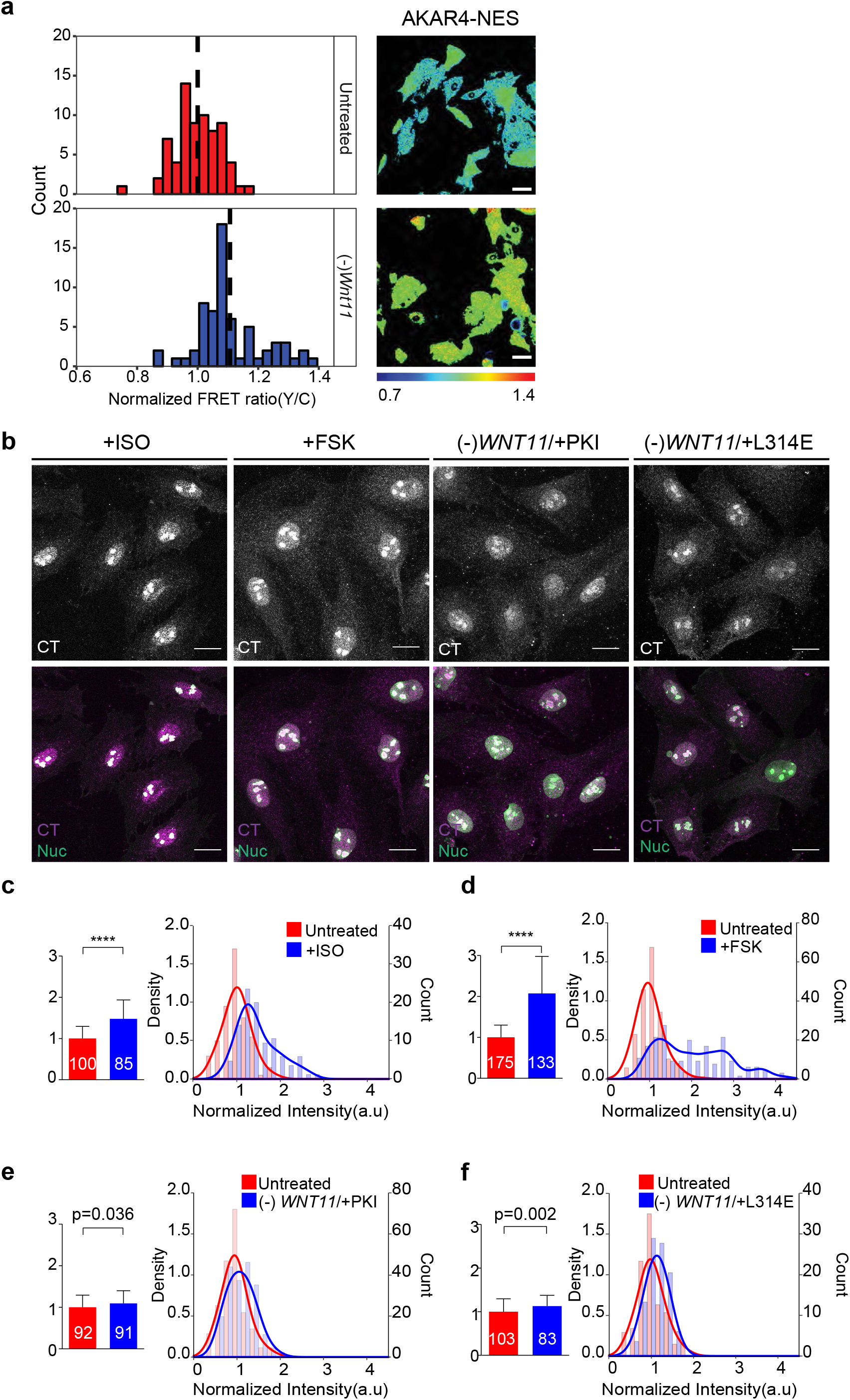
Wnt11 signaling regulates the CT formation via AKAP anchored PKA signaling. **a** Measurements of basal PKA activities using the PKA biosensor AKAR4-NES in untreated (top) and *WNT11* siRNA-treated (bottom) H9c2 cells. Normalized FRET ratios calculated from >100 frames and N ≥ 3 experiments. Dashed-line indicates means. Unpaired t-test with Welch’s correction; \*\*\*\**P*<0.0001. Scale bar, 50 μm. **b** Representative maximum intensity Z-projection of CT signals in H9c2 cells treated with isoprotenerol (ISO, 4μM), Forskolin (FSK, 10μM), *WNT11* siRNA together with protein kinase inhibitor (PKI, 10μM), or L314E peptide (100μM) (gray scale, upper panels) and merged (magenta, lower panels) with nucleolin (Nuc) signals (green). Scale bar, 20 μm. **c-f** Quantification of mean intensity of CT signals in nucleoli shown in (**b**). Bar graphs show mean ± SD of N ≥ 3 experiments and n ≥ 83 cells. Unpaired t-test with Welch’s correction; \*\*\*\**P*<0.0001. Corresponding histograms overlaid with density plots show the distribution of normalized pixel intensity values within all analyzed images.

To achieve specificity of PKA signaling, the kinase is often tethered to its substrates by AKAPs^13–15, 40^. To test whether the observed effect of PKA activation involves its interaction with AKAPs, we globally uncoupled the kinase from AKAPs using the peptide AKAP18δ-L314E^41^. This peptide binds the regulatory subunits of PKA with nanomolar affinity, and effectively blocks AKAP-PKA interaction^41^. We used the membrane-permeant version of the peptide (L314E) to confirm that PKA requires AKAP-anchoring to phosphorylate its substrates. The peptide mildly reduced the levels of some phosphorylated PKA substrates (e.g. 250 kDa) (Supplementary Fig. 2b), but importantly it reverted the CT formation in the absence of *WNT11* (Fig. 2a, e). Thus, our data indicated Wnt11 prevents the CT generation via AKAP-anchored PKA signaling.

To identify which AKAP(s) might have a role in the CT isoform formation downstream of Wnt11 signaling, we screened potential AKAP candidates (Fig. 3a). The AKAP protein family comprises 17 annotated human *AKAP* genes, which encode as many as 89 protein-coding transcripts (Supplementary Table 1). From available databases, we selected 13 that are expressed in the human heart. Out of these 13, 7 AKAP proteins showed expression or significantly differential expression in differentiated vs. non-differentiated C2C12 mouse myoblast cells (Fig. 3b). On the basis of three Gene Ontology terms (i.e. membrane localization, PKA binding domain, and ion-channel binding domain), we filtered out 6 AKAPs (Fig. 3a). As we sought to identify the conserved role of identified AKAP(s) in Wnt11 signaling, a further criterion was that the selected AKAPs must have a zebrafish orthologue, leaving us with the following list: *AKAP2*, *AKAP6*, *AKAP10*, *AKAP11*, *AKAP12*, and *AKAP13* (Fig. 3a). The zebrafish orthologues of these genes are: *palm2*, *akap6*, *akap10*, *akap11*, *akap12*, and *akap13*, respectively. We extended this list with *AKAP5* and *AKAP7*, since many studies demonstrated their role in the LTCC regulation^17–20^, although *akap5* does not have a zebrafish orthologue. Of note, *palm2* encodes a read-through transcript of Palm2 and Akap2 encoded by the *palm2-203* and *palm2-201/202*, respectively, conserved in at least human and mouse. Palm2 and Akap2 are unrelated proteins with different function. We refer to zebrafish *palm2*-*201/202* as *akap2* in this study.

**Fig. 3.**
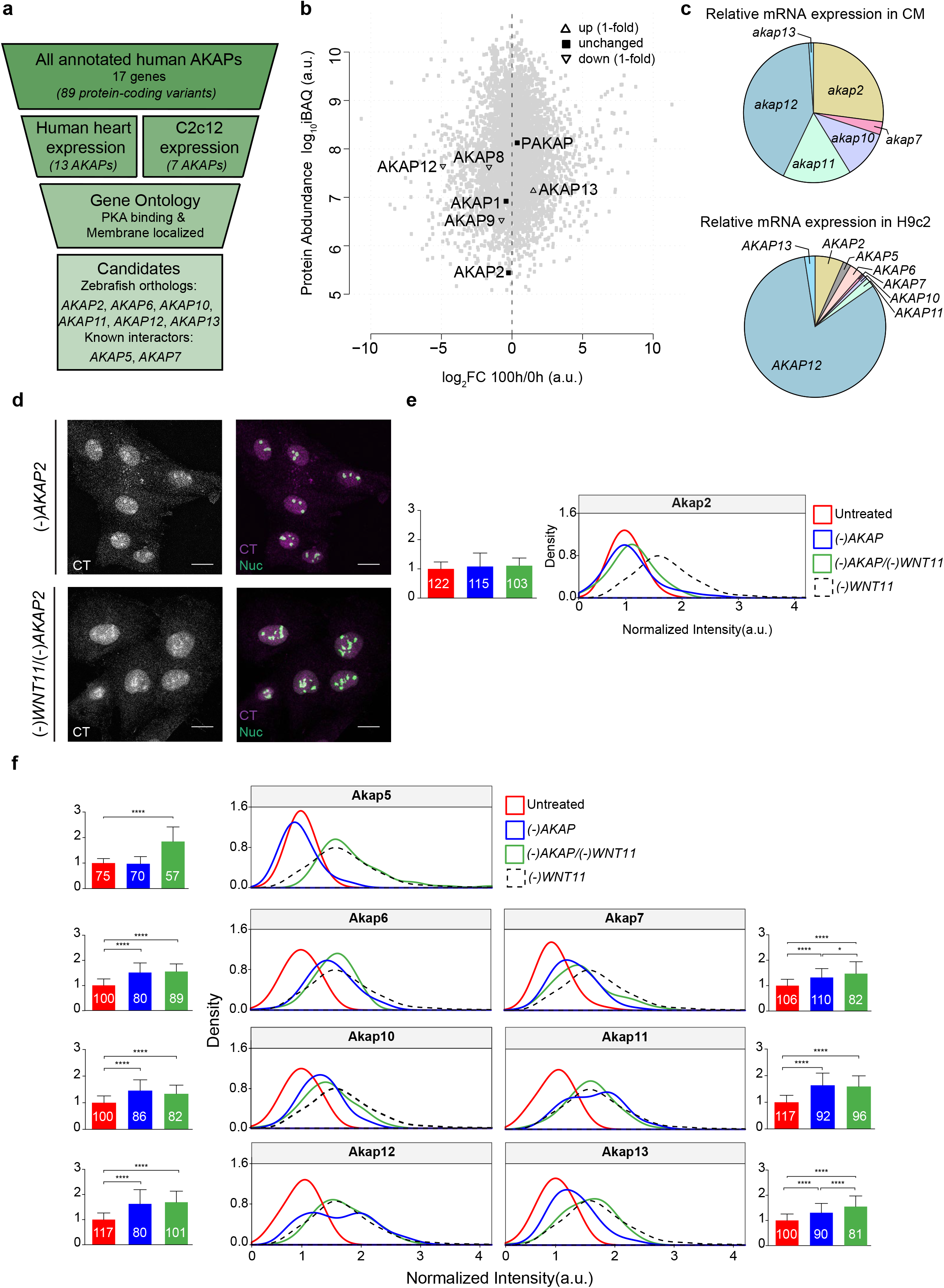
AKAP2-anchored PKA signaling regulates the CT formation downstream of Wnt11. **a** Schematic of AKAPs candidate mini-screen. **b** Expression profile of AKAPs in C2C12 cells during differentiation. The log_2_ fold change of the 100h (log_2_-FC) and 0h timepoint were used to identify changes in protein abundance between myotubes (100h) and myoblasts (0h). Log-FC were plotted against relative protein abundance estimates (iBAQ). **c** Relative mRNA expression of the selected *AKAP*s in FACS-sorted cardiomyocytes from *Tg(myl7*:*EGFP)* normalized to *myl7* (top), and in H9c2 cells normalized to *GAPDH (bottom)* (N= 3 experiments). **d** Representative maximum intensity Z-projection of CT signals in *AKAP2* siRNA-transfected H9c2 cells alone, or in combination with *WNT11* siRNA (gray scale, right panels) and merged (magenta, left panel) with nucleolin (Nuc) signals (green). Scale bar, 20 μm. **e** Quantified mean intensity of CT signals in nucleoli of untreated control (red) and *AKAP2* siRNA-transfected H9c2 cells alone (blue), or in combination with *WNT11* siRNA (green). Bar graphs show mean ± SD of N ≥ 3 experiments. Differences compared to untreated control were analyzed by One-way ANOVA with Tukey’s multiple comparison test, *P*=0.06. Number in each column indicates cell number (n). Corresponding density plots show the distribution of normalized pixel intensity values within all analyzed images. Distribution of signals in *(-)WNT11* cells are shown as reference (dotted black line). **f** Quantification of CT signals in nucleoli, both in single knockdown of remaining AKAP candidates and double knockdown experiments with *WNT11* siRNA, as described in (B). Bar graphs show mean ± SD of N ≥ 3 experiments. One-way ANOVA with Tukey’s multiple comparison test, \**P*<0.05, \*\*\*\**P*<0.0001. Number in each column indicates cell number (n).

To determine whether these genes are expressed in embryonic zebrafish heart, we used the *Tg(myl7:eGFP)*^twu3442^, in which the minimal promoter of *myl7* (*myosin, light chain 7, regulatory*) drives the heart-specific expression of *eGFP*. We performed FACS (Fluorescence-activated cell sorting) on 54 hours post fertilization (hpf) old embryos, followed by qPCR, and quantified the expression level of the aforementioned *akaps* in sorted cardiomyocytes (CM) compared to non-CM. We found that *akap6* is only expressed in non-CM, while *akap11* is only expressed in CM. *Akap2*, *akap7*, and *akap12* showed significantly higher expression levels in CM (Supplemntary Fig. 2c). Next, we sought to identify the relationship of relative mRNA expression levels between the different *akaps* in CM (Fig. 3c). Our data show that *akap12* is present in the highest amount with 41.8% followed by *akap2* with 27.1% (Fig. 3c). Corroborating these results, we performed similar quantification in H9c2 cells (Fig. 3c). mRNA expression levels of selected *AKAPs* in H9c2 cells revealed a similar expression pattern: *AKAP12* being the most abundant with 82.3% followed by *AKAP2* with 6.6% (Fig. 3c).

### AKAP2/PKA signaling regulates the CT formation downstream of Wnt11

To identify which of the selected AKAPs transduces the Wnt11 signal in preventing the CT formation, and ultimately results in LTCC regulation, we knocked down these *AKAPs* individually, or together with *WNT11* by siRNA treatment, as confirmed by qPCR (Supplementary Fig. 3a-h). Immunostaining revealed the loss of all but *AKAP2* and *AKAP5* increased CT isoform formation (Fig. 3d-f, Supplementary Fig. 3i-o). Markedly, only the loss of *AKAP2* could revert the loss of *WNT11* phenotype in a similar fashion as previously observed with PKI or L314E treatment (compare Fig. 3d, e to Fig. 2b, e, f). These results indicated a functional redundancy between CM-specific AKAPs in generating the CT isoform. Most importantly, our data demonstrated that Wnt11 signaling specifically targets AKAP2 to compartmentalize PKA activity as a mechanism to attenuate LTCC activity.

### AKAP2 physically interacts with LTCC

Next, we investigated whether the AKAP2-dependent regulation requires complex formation with the LTCC. As a means to determine whether AKAP2 interacts with the CT, we designed a construct encoding the rat CT from amino acid (aa) 1342 to 2006 tagged with FLAG on the N terminus and mVenus on the C terminus as illustrated in Fig. 4a. To show that we can efficiently immuno-precipitate (IP) this construct, we overexpressed it in H9c2 cell, immunoprecipitated the protein through its tags, and detected the precipitated proteins by Western blotting (Fig. 4b, Supplementary Fig. 4a). Detection of the proteins precipitated via the FLAG tag with FLAG antibody identified 3 isoforms: an isoform of around 130 kDa, which corresponds to the full-length construct, a shorter 80 kDa fragment, which corresponds to the CT construct without mVenus, and a 30 kDa fragment, which corresponds to the pCT. Although the predicted full-length CT construct is 105 kDa, these data indicate that its mobility on SDS-PAGE is 130 kDa. In addition, we identified 7 extra bands, which may be explained by the existence of multiple cleavage sites, or other splice variants of the CT construct, or the combination of both. By probing the FLAG precipitate with a GFP antibody, we detected the full-length 130 kDa isoform. Precipitation through GFP followed by the detection with anti-FLAG antibody also revealed a 130 kDa isoform. However, detection of the GFP-IP with anti-GFP antibody identified two other isoforms in addition to a 130 kDa isoform: a 90 kDa one corresponding to dCT-mVenus, and a 30 kDa one, which is most likely the cleaved mVenus alone. Altogether, the existence of a 30 kDa isoform of FLAG-IP/FLAG detection, and a 90 kDa isoform of GFP-IP/GFP detection indicate our exogenously expressed CT construct is proteolytically processed similarly to the endogenous LTCC at the calpain cleavage site^11^.

**Fig. 4.**
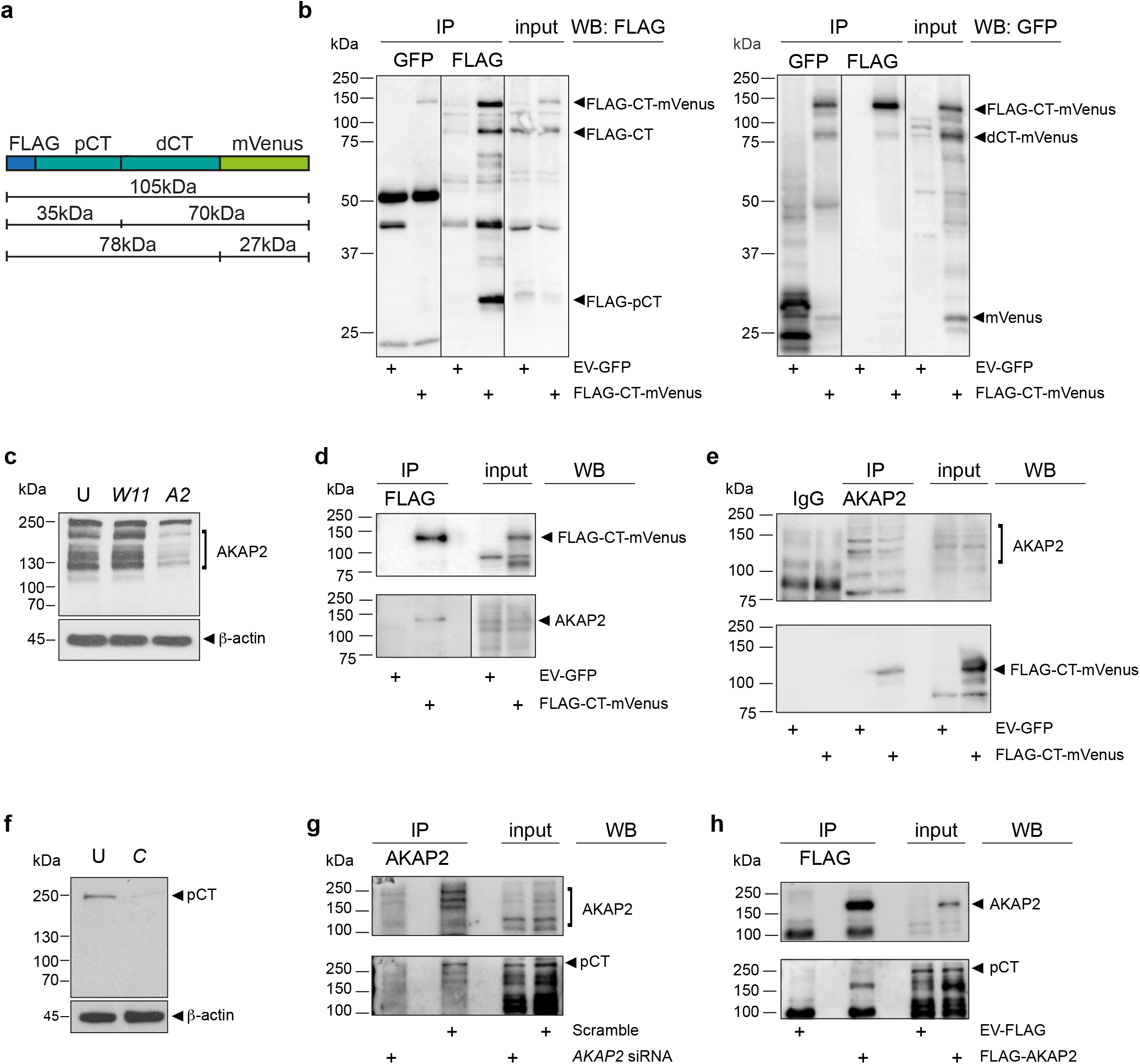
AKAP2 directly binds to LTCC. **a** Schematic of the FLAG-and mVenus-tagged CT construct (UniProt ID: A0A0G2QC25, 1342aa-2006aa) and the expected CT isoforms after proteolytic processing. **b** FLAG (left panel) or GFP (right panel) detection by WB of CT construct of lysates of H9c2 cells, transfected either with the CT construct or with empty vector GFP (EV-GFP) control, showing input and FLAG-or GFP-IP. N ≥ 10 experiments. **c** Western blot (WB) detection of AKAP2 in lysates of untreated control (U) and *WNT11* (*W11*) and *AKAP2* (*A2*) siRNA transfected H9c2 cells by using anti-AKAP2 antibody. Membrane was stained with β-actin as loading control. (N = ≥ 5 experiments). **d** GFP (upper panel) and AKAP (lower panel) Co-IP detection by WB of lysates of H9c2 cells transfected either with the CT construct or EV-GFP control, showing input and FLAG-IP. N ≥ 3 experiments. **e** AKAP2 (upper panel) and FLAG (lower panel) Co-IP detection by WB of lysates of H9c2 cells transfected either with the CT construct or EV-GFP control, showing input and endogenous AKAP2-IP. N = 3 experiments. **f** CT detection by WB in lysates of untreated control (U) and *CACNA1C* siRNA-transfected (C) H9c2 cells using anti-pCt antibody. Membrane was stained with β-actin as loading control. (N = ≥ 3 experiments). **g** AKAP2 (upper panel) and pCT (lower panel) Co-IP detection by WB of lysates of H9c2 cells transfected either with scramble or *AKAP2* siRNA, showing input and endogenous AKAP2-IP. N = 1 experiments. **h** AKAP2 (upper panel) and pCT (lower panel) Co-IP detection by WB of lysates of H9c2 cells transfected either with FLAG-AKAP2 or EV-GFP control, showing input and FLAG-IP. N = 1 experiments.

We next probed the physical interaction between the CT construct and AKAP2. AKAP2 is expressed in at least 6 isoforms^43^. Consistently, immunoblotting of H9c2 lysates identified at least 5 AKAP2 isoforms with mobility from 100 to 180 kDa (Fig. 4c). FLAG-IP of the CT construct followed by the detection with anti-AKAP2 antibody yielded a band of around 130 kDa (Fig. 4d, Supplementary Fig. 4b) corresponding to one of the AKAP2 isoforms. Reciprocal co-IP of the endogenous AKAP2-IP followed by the detection of the CT construct using its FLAG tag (Fig. 4e, Supplementary Fig. 4c) validated our results, and demonstrating a physical interaction between the CT and AKAP2.

To study the interaction between endogenous proteins, we generated a custom-made antibody (rat anti-pCt), which recognizes the CT at aa 1758-1777. We first verified the specificity of this antibody (Fig. 4f). The anti-pCt antibody recognizes a 250 kDa band corresponding to the full-length α1C subunit. After AKAP2-IP/AKAP2 detection, we re-stained the membrane with the anti-pCt, which resulted in an additional signal at around 250 kDa in Scramble siRNA-treated, but not in *AKAP2* siRNA-treated cells (Fig. 4g, Supplementary Fig. 4d). This 250 kDa band corresponds to the full-length α1C subunit, indicating an interaction between AKAP2 and the channel. To confirm this interaction, we overexpressed an AKAP2 construct, which is tagged with FLAG and HA on its N terminus. After AKAP2-IP followed by AKAP2 detection, we re-stained the membrane with our custom-made pCT antibody. In cells transfected with the tagged AKAP2 construct, an additional band appeared around 250 kDa, but not in the cells transfected with an empty control plasmid (Fig. 4h, Supplementary Fig. 4e). In summary, these data provided strong evidence that AKAP2 interacts with the LTCC, either directly or via a yet unidentified complex, and coordinates local Wnt11 signaling.

### Akap2 is essential for heart development in zebrafish

To gain insight into Akap2 function during cardiac development, we utilized the developing zebrafish embryo as the *in vivo* model. We showed by qPCR that *akap2* is highly expressed in embryonic CM (Fig. 3c, Supplementary Fig. 2c). Because *akap2* expression pattern within the whole embryo is unknown, we performed *in situ* hybridization (ISH). In wild-type (WT) embryos, we detected strong *akap2* expression in the brain, the eyes, and in the heart at 48 hpf (Fig. 5a, Supplementary Fig. 5a). Using complementary approaches, we characterized the consequences of the loss of *akap2* (Fig. 5b-f, Supplementary Fig. 5b-h). Knock down of *akap2* by using a splicing-site targeting morpholino (*akap2*^e2i2^ MO) resulted in the exclusion of exon 2, which includes the PKA binding domain (*akap2Δex2*) (Fig. 5b). RNA extraction followed by RT-PCR confirmed that only *akap2*^e2i2^-, but not control-or mismatch MO-injected embryos, lacked the exon 2 of *akap2* as expected (Fig. 5c). Of these embryos, 43.4% were shorter and had smaller head and eyes compared to WT (Fig. 5d, Supplementary Fig. 5b). They also developed cardiac edema and looping defects (Fig. 5e, f). Similarly, both the injection of a morpholino targeting the transcription start site (*akap2*^ATG^ MO) and mosaic CRISPR/Cas9 somatic mutagenesis with *akap2* sgRNA targeting exon 2 (Fig. 5d, Supplementary Fig. 5c-e) yielded comparable phenotypes in 48% and 41.6% of embryos, respectively. Neither injection of control or mismatch morpholino, nor injection of Cas9 alone induced the observed phenotypes (Fig. 5d, Supplementary Fig. 5f, h).

**Fig. 5.**
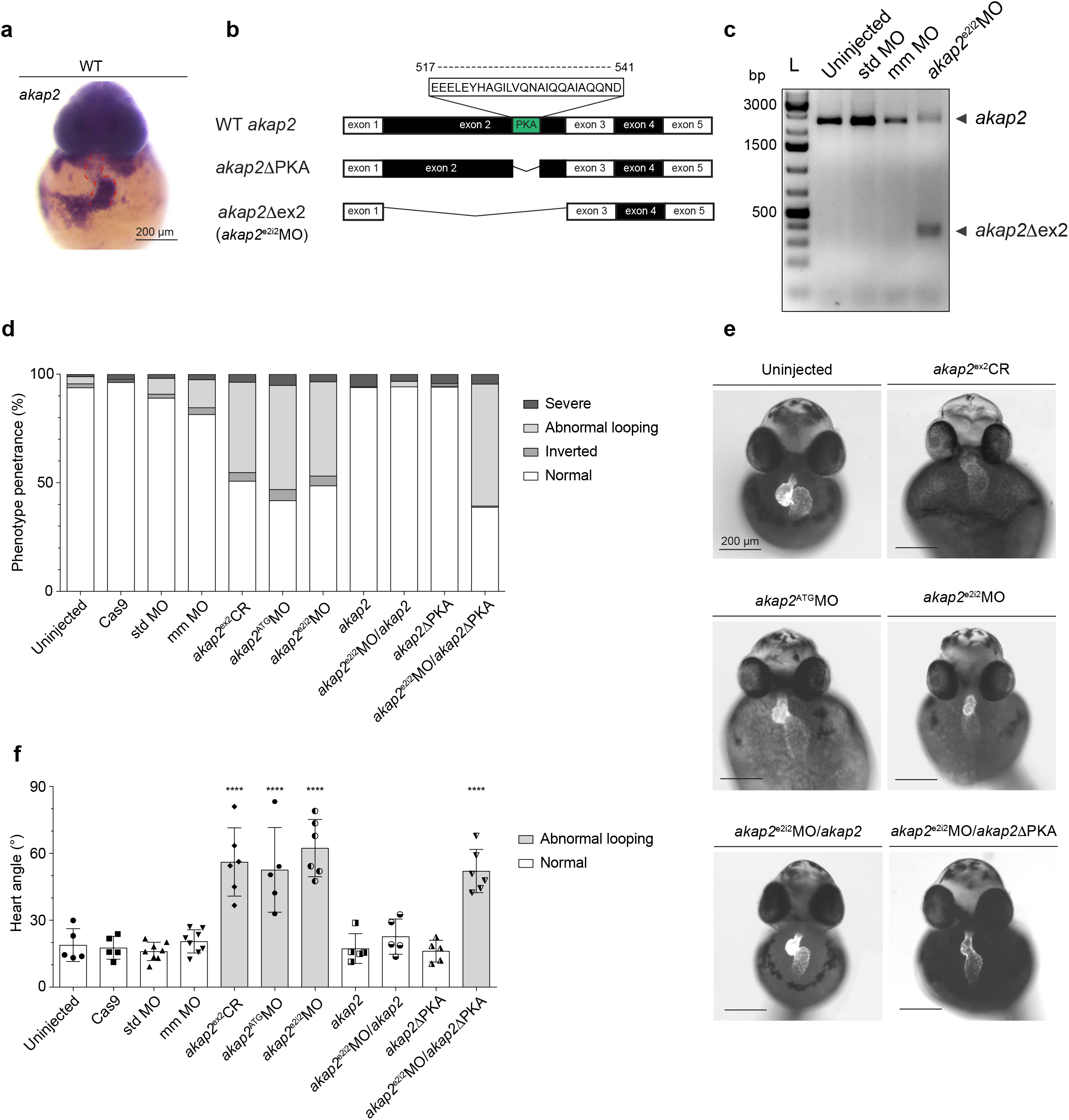
Akap2 is essential for normal heart development in zebrafish. **a** *In situ* hybridization detection of endogenous *akap2* mRNA in whole-mount wild-type (WT) zebrafish embryo at 48 hpf. Frontal view of an embryo shows head- and heart-specific (red dashed line) expression pattern of the *akap2*. Scale bar, 200 μm. **b** Schematic of the zebrafish wild-type (WT) *akap2* (palm2-202 ENSDART00000165942.2) with predicted PKA binding domain (green), as well as of *akap2*ΔPKA mRNA lacking PKA binding domain, and *akap2*^e2i2^ morpholino-(MO) induced exclusion of exon 2 (*akap2*Δex2). The exons are not depicted to scale. PKA: Protein-kinase-A binding domain. **c** RT-PCR of *akap2* cDNA from uninjected control, standard morpholino-injected (std MO), mismatch morpholino-injected (mm MO) and *akap2*^e2i2^MO-injected embryos at 54hpf. The size of the *akap2* fragment is 2434 bp, while in *akap2* morphants it is only 300 bp due to exclusion of the exon 2 (akap2Δex2). **d** Phenotypic analyses of the loss of *akap2* based on heart defects. N ≥ 3 experiments; n ≥ 300 embryos. **e** Bright field image overlayed with fluorescent image, frontal view of uninjected control, *akap2* sgRNA/Cas9-injected (*akap2*^ex2^CR), *akap2* morpholino-(*akap2*^e2i2^MO or *akap2*^ATG^MO) injected, *akap2*^e2i2^MO with WT *akap2-* or *akap2*ΔPKA-mRNA injected *Tg(myl7*:*EGFP)* zebrafish embryos at 54hpf. Scale bar, 200 μm. **f** Quantification of the heart-looping angle. Data show mean ± SD of at least 5 embryos per condition. One-way ANOVA with Dunnett’s multiple comparison test, *P*_Cas9_=0.999; *P*_std MO_=0.999; *P*_mm MO_=0.999; *P_akap2_*_mRNA_=0.999; *P_akap2_*_e2i2MO/*akap2* mRNA_=0.996; *P_akap2_*_ΔPKA mRNA_=0.9993. **** *P*<0.0001.

In developing zebrafish embryos, cardiac looping occurs from 30 to 48 hpf when the linear heart tube becomes bicameral, and the ventricle and atrium become morphologically distinguishable. To visualize the looping defect, we injected *Tg(myl7:eGFP)* with either aforementioned morpholinos, or an *akap2* sgRNA/Cas9 nucleoprotein complex and measured the heart looping angle at 54 hpf (Fig. 5e, f). The defined angle is within the midsagittal line and the atrioventricular canal, and indicates the relative position of the atrium to the ventricle. In 54 hpf uninjected control embryos, the mean looping angle was 18.9° ± 7.4°, with the atrium in an almost juxtaposed position to the ventricle (Fig. 5e, f). On the other hand, injection of either *akap2* sgRNA, or *akap2*^ATG^ and *akap2*^e2i2^ morpholinos increased the looping angle to 56.1° ± 15.3°, 52.6° ± 19.0°, and 62.4° ± 12.8° respectively (Fig. 5e, f). Injection of control or mismatch morpholino or injection of Cas9 alone did not induce any phenotypes (Fig. 5f, Supplementary Fig. 5h).

To ensure these phenotypic characteristics outlined above are caused by the loss of *akap2* and are not off-target effects of the morpholino, we cloned *akap2* mRNA and performed rescue experiments. Injection of *akap2* mRNA alone did not cause any defects (Fig. 5d, Supplementary Fig. 5b), and the heart looping was comparable to WT at 17.3° ± 6.7° (Fig. 5f, Supplementary Fig. 5h). Co-injection of *akap2* mRNA and *akap2*^e2i2^ MO completely abolished the morpholino effect and resulted in normally developed embryos without any visible phenotype with a mean looping angle of 22.7° ± 7.9° (Fig. 5d-f, Supplementary Fig. 5b). To further verify these results, we cloned *akap2* mRNA without the PKA binding domain (*akap2ΔPKA*) (Fig. 5b). We surmised this should abolish AKAP2-specific compartmentalization of PKA activity, which our *in vitro* data have suggested to be a central mechanism of Wnt11-dependent LTCC regulation. Indeed, co-injection of *akap2ΔPKA* mRNA with *akap2*^e2i2^ MO failed to rescue the morpholino-induced phenotype with 56.2% affected embryos, and an average looping angle of 48.9° ± 6.6° (Fig. 5d-f, Supplementary Fig. 5g), while injection of *akap2ΔPKA* mRNA alone yielded normal embryos with the looping angle of 16.2° ± 4.9° which is comparable to WT (Fig. 5d, f, Supplementary Fig. 5g, h). Taken together, our data showed AKAP2, specifically through its scaffolding function for PKA, is essential for cardiac development.

### AKAP2 regulates the intracellular calcium concentrations

Our findings prompted us to examine whether Akap2 affects the phasic changes of intracellular calcium concentrations [Ca^2+^]_i_ during diastole and systole similarly to Wnt11. Using high-speed ratiometric calcium imaging, we measured calcium transients in WT and *akap2*-deficient zebrafish hearts as previously reported^32^. Loss of *akap2* markedly increased calcium transient amplitudes when compared to WT hearts (Fig. 6a-c). Significant increases were readily observed in the atria, with larger variability in the ventricles during systole (Fig. 6b, c), while the diastolic [Ca^2+^]_i_ seemed unaffected, in either the atrium or in the ventricle (Fig. 6d).

**Fig. 6.**
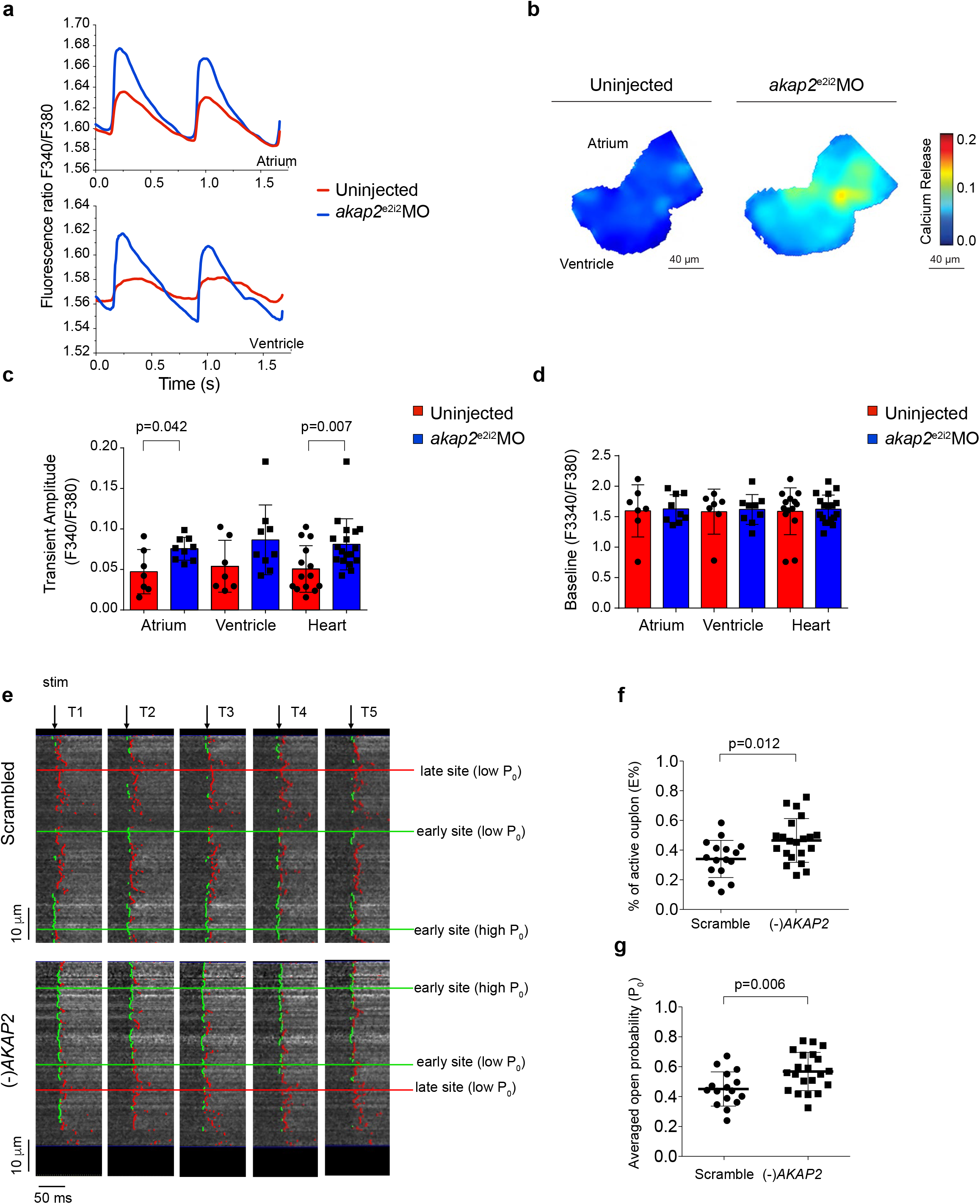
AKAP2 regulates the LTCC conductance. **a** Averaged Ca^2+^ transients from uninjected control (red) and *akap2*^e2i2^MO-injected (blue) zebrafish embryonic hearts at 54 hpf in atrium (upper panel) and ventricle (lower panel). **b** Ratiometric map of Ca^2+^ transient amplitudes from uninjected control and *akap2*^e2i2^MO-injected zebrafish embryonic hearts at 54 hpf. Scale bar, 40 μm. **c** Bar graphs show mean ± SD of Ca^2+^ transient amplitudes in atrium, ventricle, and whole heart from uninjected control (red) and *akap2*^e2i2^MO-injected (blue) zebrafish embryos at 54 hpf. Mann-Whitney test, *P*_Ventricle_=0.071; N=3 experiments, n (*akap2*^e2i2^MO-injected) = 9, and n (uninjected control) = 7. **d** Bar graphs show mean ± SD of baseline in atrium, ventricle, and whole heart from uninjected control (red) and *akap2*^e2i2^MO injected (blue) zebrafish embryos at 54 hpf. Mann-Whitney test, *P*_Atrium_=0.837, *P*_Ventricle_=0.607, *P*_Heart_=0.64; N=3 experiments, n (*akap2*^e2i2^MO-injected) = 9 and n (uninjected control) = 7. **e** Line scan images of 5 consecutive calcium transients (Fluo4-AM, field stim, 1 Hz) in a representative NRVM. Black arrow indicates time of electrical stimuli. The green and red vertical line in each transient indicates the relative position of the time of half maximal calcium release at that site. Green is < 6 ms after begin of transient (early release), red is ≥ 6 ms as late release (same cut-off for all cells/groups). 10 consecutive transients from each cell were assessed (only 5 are shown for space reasons). **f-g** Measurements of calcium release at LTCC-RyR couplons in *AKAP2* siRNA-transfected NRVM compared to scramble control. Data express the percentage of line scanned with active couplons (**f**). Data represent the average probability of calcium releases at the couplons (**g**). Mean ± SD of N = 3 experiments, and n ≥ 16 cells. Unpaired t-test.

To explore whether the function of AKAP2 in regulating [Ca^2+^]_i_ is conserved, and to investigate more directly whether AKAP2 affects LTCC conductance, we probed the changes in calcium release in NRVM. We compared the percentage (E%) of active LTCC-RyR couplons as well as their average open probability (P_0_), which directly relate to the calcium release^33^, in ten consecutive Ca^2+^ transients (Fig. 6e). Both the active percentage and open probability of LTCC-RyR couplons are significantly increased in the absence of *AKAP2* (Fig. 6f, g). Altogether, these data highlighted AKAP2 as an integral component of the machinery regulating LTCC conductance in cardiomyoytes.

### Akap2 is an effector of Wnt11 signaling required for cardiomyocyte intercellular coupling

Wnt11/LTCC signaling regulates the emergence of the electrical coupling gradient in the developing myocardium^32^. We assessed whether Akap2 is the effector through which Wnt11 attenuates LTCC to modulate the intercellular coupling across the myocardium. We performed genetic epistasis to determine the extent of the interaction between Akap2 and Wnt11 by co-injecting *akap2*^e2i2^ and *wnt11*^ATG^ morpholino into one-cell stage zebrafish embryos (Fig. 7a, b, Supplementary Fig. 6a-c). In double morphants, about 60% of the embryos showed heart edema, abnormal looping, and cyclopic eyes—the phenotypic hallmarks of *wnt11* morphants and mutants^44^ (Fig. 7a, Supplementary Fig. 6a, b). To ensure that the embryos used for further studies were deficient for both *wnt11* and *akap2*, we selected only embryos with cyclopic eyes. RT-PCR confirmed the exclusion of *akap2* exon 2 in these embryos (Supplementary Fig. 6c). Focusing on the looping defect, loss of *wnt11* resulted in the defective looping with a heart angle of 32.5° ± 13.1°. In double *wnt11; akap2* morphants, this angle was further increased to 47.9° ± 3.1° (Fig. 7a, b). These data suggested Akap2 may genetically interact with Wnt11.

**Fig. 7.**
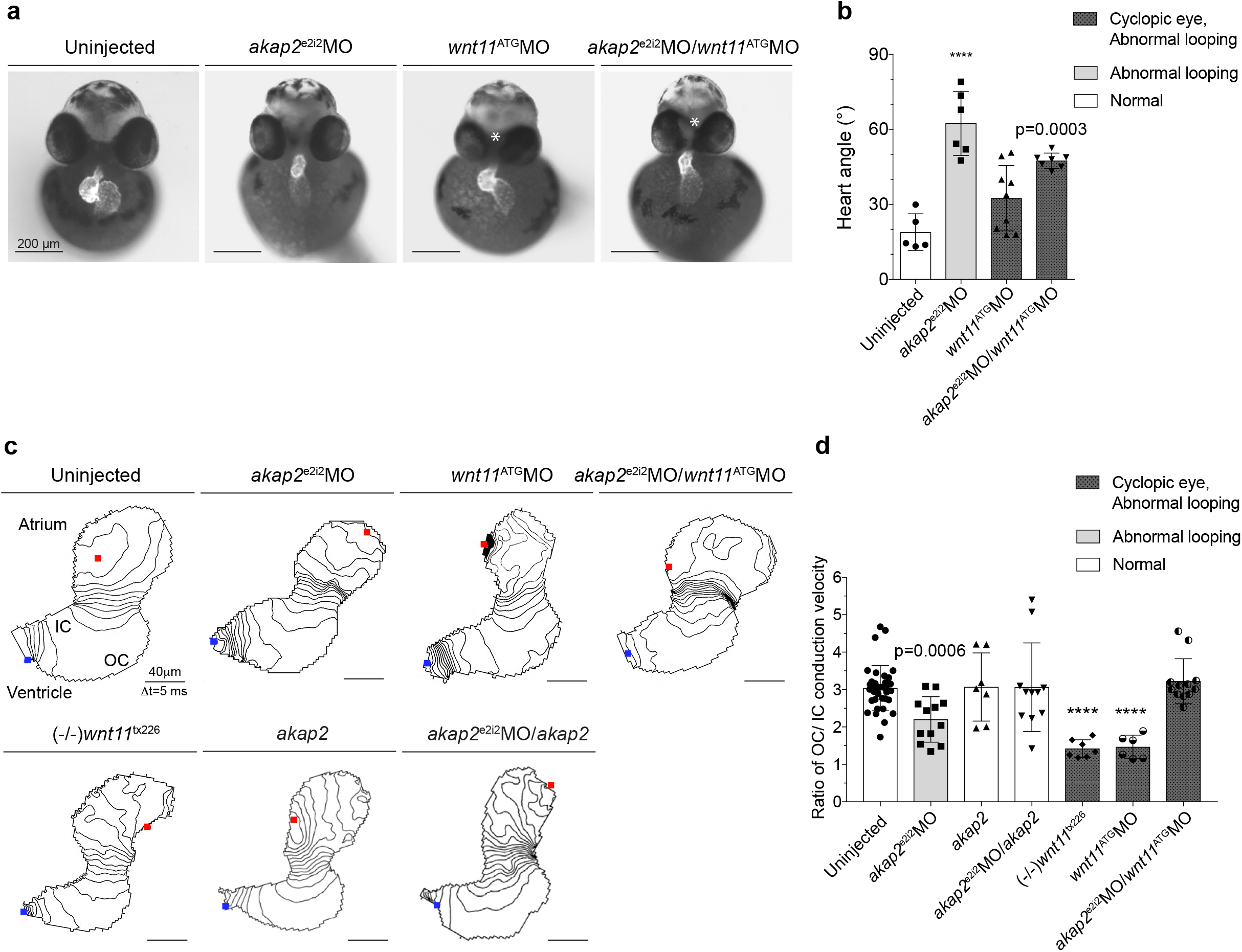
Wnt11/Akap2/PKA signalosome is required for the electrical gradient formation in developing heart. **a** Bright field image overlayed with fluorescent image, frontal view of uninjected control, *akap2*^e2i2^MO-injected, *wnt11*^ATG^MO-injected and double morpholino-injected *Tg(myl7*:*EGFP)* zebrafish embryos at 54hpf. Scale bar, 200 μm. White asterisk labels cyclopic eye, a hallmark of loss of *wnt11* phenotype. **b** Quantification of the heart-looping angle. Data show mean ± SD of at least 5 embryos per condition. One-way ANOVA with Dunnett’s multiple comparison test, *P*(*wnt11*^ATG^MO)=0.069; **** *P*<0.0001. **c** Isochronal map of 54 hpf embryonic zebrafish heart from uninjected control, *akap2*^e2i2^MO- or *wnt11*^ATG^MO- or *akap2*^e2i2^MO/*wnt11*^ATG^MO-injected embryos, from *wnt11* mutant ((-/-)*wnt11*^tx226^), from *akap2* mRNA-injected (*akap2*) or *akap2* mRNA-and *akap2*^e2i2^MO-injected embryos. Lines represent the positions of the action potential wavefront at 5 ms intervals. Red dots indicate the start site of the excitation and blue dots represent the end. Scale bar, 40 μm. IC, inner curvature, OC, outer curvature. **d** Quantification of the voltage propagation at 54 hpf of uninjected control, *akap2*^e2i2^MO-or *wnt11*^ATG^MO-or *akap2*^e2i2^MO/*wnt11*^ATG^MO-injected embryos, from *wnt11* mutant ((-/-)*wnt11*^tx226^), from *akap2* mRNA-injected (*akap2*) or *akap2* mRNA- and *akap2*^e2i2^MO-injected embryos. Data express the ratio of the OC/IC conduction velocity and are plotted as mean ± SD of N=3 experiments. One-way ANOVA with Dunnett’s multiple comparison test, *P*(*akap2*)=0.999; *P*(*akap2*^e2i2^MO*/akap2*)=0.145, *P*(*akap2*^e2i2^MO*/wnt11*^ATG^MO)=0.985, *\*\**\*\**P*<0.0001.

The action potential (AP) triggered by the sinus node propagates from the atrium through the atrioventricular junction to the ventricle, where in normal conditions at 54 hpf the outer curvature (OC) of the ventricle conducts AP three times faster than the inner curvature (IC)^32^ (Fig. 7c, d). To examine the physiological role of Akap2 in AP propagation, we performed high-speed optical mapping of transmembrane potentials in the zebrafish heart (Fig. 7c, d). We measured the conduction velocities in the OC and IC regions of interest and plotted them as an OC/IC ratio. We confirmed in uninjected controls this ratio was close to three (3.21 ± 0.54). In *wnt11*-deficient hearts, the gradient formation was completely abolished with a ratio of 1.47 ± 0,32, and 1.65 ± 0.70 in *wnt11^tx2^*^26^ mutants (Fig. 7c, d) as previously reported^32^. The OC/IC ratio was significantly reduced in the absence of *akap2* (2.20 ± 0.61) (Fig. 7c, d) indicating that Akap2 contributes to the patterning of intercellular electrical coupling. Co-injection of *akap2* mRNA with *akap2*^e2i2^ MO yielded a ratio of 3.06 ± 1.18, thus restoring the WT coupling gradient (Fig. 7c, d). Injection of *akap2* mRNA alone led to unaltered intercellular coupling with a ratio of 3.07 ± 0.91 (Fig. 7c, d). Importantly, in the hearts deficient for both *wnt11* and *akap2*, the formation of the intercellular electrical gradient was completely restored, with a ratio of 3.22 ± 0.60 (Fig. 7c, d), further demonstrating that Akap2 is an integral component of Wnt11/LTCC signaling.

Taken together, we have identified AKAP2 as a downstream effector of the WNT11/LTCC pathway, where compartmentalization of AKAP2-anchored PKA signaling is required for tuning LTCC conductance.

## Discussion

Stimulation via the β-adrenergic signaling cascade and shifting the L-type calcium channel gating mode to more open probability states has been thoroughly studied as a signaling pathway regulating the channel conductance^5,7^. Lesser attention, however, has been given to mechanisms independent of classical GPCR systems that attenuate the ionic influx through the LTCC. Here, we describe a novel branch of Wnt11 non-canonical signal transduction: Wnt11 together with its receptor Fzd7 compartmentalize PKA activity scaffolded via a complex with AKAP2. We demonstrate a previously uncharacterized interaction between AKAP2 and the C-terminus of the LTCC, the result of which modulates channel conductance. Wnt11 signaling prevents this interaction, thus lowering the channel open probability states. In heart development and physiology, this Wnt11/AKAP2/PKA-dependent attenuation of LTCC conductance is crucial for the emergence of the intercellular electrical coupling gradients necessary for the formation of sequential cardiac contraction.

Frizzled proteins are Wnt ligand receptors belonging to the Class F receptors of GPCR family^24^. The unique sequence and structural features of the C-terminal tail of Frizzleds are noted to hinder the binding of traditional GPCR ligands^24, 26^. Nevertheless, increasing evidence are indicative of their ability to signal through heterotrimeric G proteins^25, 27, 45^. Similarly, Wnt ligands are able to regulate cAMP levels and thus activate the cAMP/PKA pathway in diverse contexts^46–48^. Our findings demonstrate that in addition, Wnt signaling can also modulate the spatiotemporal regulation of PKA activation. In constrast to the β-adrenergic system, we propose Wnt signaling prevents PKA-dependent phosphorylation of its targets, in this case the C-terminal tail of the LTCC, by compartmentalizing AKAP/PKA complexes. Compartmentalization of AKAP/PKA signalosomes, which is essential for PKA substrates recognition and PKA activity^40, 49^, can provide specificity in targeting different effectors of the conserved Wnt/Fzd G protein-coupled signaling.

In cardiomyocytes, a number of AKAPs have been identified^50^. Thus far, only AKAP5 (AKAP79/150) and AKAP7 (AKAP15/AKAP18) of the AKAP family have been described in regulating the LTCC^18, 20, 21, 51^. Here we refine the catalog of cardiomyocyte-specific AKAPs with an emphasis on those which are developmentally regulated. Of note, although there are other AKAP-domain containing proteins binding and scaffolding PKA, either in the Wnt pathway^52^ or implicated in the LTCC regulation^53^, we chose to focus in this study on the bonafide AKAP protein family consisting of 17 members (Supplementary Table 1). Out of 6 potential AKAPs, we found only AKAP2 transducing Wnt11/Fzd7 signals to regulate the Ca^2+^ influx through the LTCC. By demonstrating the physical binding of AKAP2 to the C-terminal tail of LTCC, we establish a novel interacting AKAP capable of regulating LTCC conductance in a PKA-dependent manner. Moreover, we establish this is a conserved interaction regulated by the Wnt11/Fzd7 signaling. Lack of Wnt11/Fzd7 induced the PKA/AKAP2-dependent cleavage of the C-terminus, leading to increased Ca^2+^ influx through the LTCC across species. Given that AKAP2 associates with the plasma membrane or the actin cytoskeleton^43, 54, 55^, it is possible Wnt11 signaling restricts AKAP2 binding to the LTCC via known effects on endocytosis or actomyosin contractlilty^44, 56–58^.

The Wnt11 ligand is indispensable for heart muscle cell specification and terminal differentiation^59–62^. Previously, we have shown that Wnt11 attenuates the LTCC conductance, which is required for proper formation of intercellular coupling and electrical gradients^32^. Loss of coupling gradients, and with them the associated physiological boundaries in the myocardial syncytium, are detrimental for the generation of physiologic contraction, and possibly for septation in higher species^63^. Besides its role in calcitonin-mediated cancer invasion^55^ and ocular transparency^54^, the cellular functions of AKAP2 are not fully explored. Mechanistically, this Wnt11/AKAP2/PKA axis attenuation of the LTCC conductance has major implications for cardiac muscle physiology, as observed by changes in calcium transient amplitudes and percentage of active LTCC-RyR couplons. Additionally, lack of *akap2* affected both heart morphology and electrical coupling. These data and prior evidence demonstrating dysregulated Wnt signaling and PKA-AKAP2 in different forms of cardiomyopathies^64–66^, imply potential links between Wnt11/AKAP2/PKA and human heart muscle diseases or arrhythmias, potentially via effects on cardiomyocyte differentiation^67^. There also exists some evidence from work in stem cells suggesting a complex role for the Wnt11/AKAP2/PKA axis in integration of electrical and mechanical inputs^68–70^. Modulation of electrical signals through Wnt11/AKAP2/LTCC signaling might therefore be important to explore in the context of stem cell differentiation in the future.

Taken together, we establish the molecular mechanism through which Wnt11 signaling attenuates the LTCC conductance. This distinct branch of non-canonical Wnt signaling represented by the multivalent effector complex of PKA/AKAP2/LTCC is required for the formation and regulation of intercellular electrical coupling in the early cardiac epithelium. Our findings reveal Wnt11/Fzd7 signaling as an alternative GPCR system to β-adrenergic cascade in modulating the LTCC conductance across species. In more general terms, we propose that the Wnt-dependent regulation of L-Type calcium channels via protein kinase compartmentalization may be a key effector in many different cell types and tissues, ranging from neurons to insulin-producing cells. Indeed, LTCC attenuation by Wnt signals may be of fundamental importance in regulating various cellular states through excitation-coupling mechanisms.

**Table 1.**

| **Gene name**<br>**(Human)** | **Other names** | **Rat**<br>**Orthologs** | **Mouse**<br>**Orthologs** | **Zebrafish**<br>**Orthologs** |
| --- | --- | --- | --- | --- |
| AKAP1 | AKAP121, AKAP149,<br>SAKAP84, S-AKAP84,<br>AKAP84, D-AKAP1,<br>PPP1R43, TDRD17 | + | + | AKAP1b |
| AKAP2<br>PALM2-AKAP2 | AKAP-KL, KIAA0920,<br>DKFZp564L0716,<br>MISP2, PKRA2 | + | + | - |
|  |  | - | + | + |
| AKAP3 | FSP95, SOB1,<br>AKAP110, CT82 | + | + | - |
| AKAP4 | p82, hAKAP82, AKAP82,<br>Fsc1, HI, CT99 | + | + | - |
| AKAP5 | AKAP75, AKAP79 | + | + | - |
| AKAP6 | KIAA0311, mAKAP,<br>AKAP100, PRKA6,<br>ADAP6 | + | + | + |
| AKAP7 | AKAP18, AKAP15 | + | + | + |
| AKAP8 | AKAP95,<br>DKFZp586B1222 | + | + | - |
| AKAP8L |  | + | + | + |
| AKAP9 | KIAA0803, AKAP350,<br>AKAP450, CG-NAP,<br>YOTIAO, HYPERION,<br>PRKA9,<br>MU-RMS-40.16A,<br>PPP1R45, LQT11 | + | + | + |
| AKAP10 | D-AKAP2, PRKA10,<br>MGC9414 | + | + | + |
| AKAP11 | KIAA0629, AKAP220,<br>PRKA11, FLJ11304,<br>DKFZp781I12161,<br>PPP1R44 | + | + | + |
| AKAP12 | AKAP250, SSeCKS | + | + | AKAP12a<br>AKAP12b |
| AKAP13 | Ht31, BRX, AKAP-Lbc, c-<br>lbc, PROTO-LB, HA-3,<br>ARHGEF13 | + | + | + |
| AKAP14 | AKAP28 | + | + | + |
| AKAP17A | XE7, XE7Y, DXYS155E,<br>MGC39904, 721P,<br>CCDC133 | + | - | + |
| AKAP17B | AKAP16B, AKAP16BP | + | + | - |

## Author Contribution

K.D.C., T.R., E.K., and D.P. designed the study. K.D.C., T.R., M.H.Q.P., M.S., P.W., K.N.M., D.Pl. A.A.W., H.Z., and M.D.S. performed experiments, and analysed the data. H.Z., M.D.S., and M.Se. performed MS analysis. K.D.C., T.R., M.H.Q.P., and D.P wrote the manuscript.

## Acknowledgments

We thank Alexander M. Meyer for technical support, the Advanced Light Microscopy team, and the Fish Facility at MDC for expert support. This work was supported by the Helmholtz Young Investigator Program VH-NG-736, Deutsche Forschungsgemeinschaft (DFG) PA2619/1-1, and Marie Curie Career Integration Grant from the European Commission (MC CIG) (WNT/CALCIUM IN HEART-322189) to D.P., and by grants from the German Centre for Cardiovascular Research (DZHK) partner site Berlin (81X210012 and B18-005 SE), the Deutsche Forschungsgemeinschaft (DFG KL1415/7-1) and the Bundesministerium für Bildung und Forschung (BMBF; 16GW0179K) to EK.

## Methods

### EXPERIMENTAL MODELS

#### Ethics statement

All zebrafish husbandry and experiments were performed in accordance with guidelines approved by the Max-Delbrück Center for Molecular Medicine at the Helmholtz Association and the local authority for animal protection (Landesamt für Gesundheit und Soziales, Berlin, Germany) for the use of laboratory animals, and followed the “Principles of Laboratory Animal Care’ (NIH publication no. 86-23, revised 1985) as well as the current version of German Law on the Protection of Animals.

#### Zebrafish husbandry

Zebrafish were maintained under continuous water flow and filtration with automatic control for a 14:10 h light/dark cycle at 28.5°C. Fertilized eggs were collected and raised under standard laboratory conditions (at 28.5°C in E3 solution (5 mM NaCl, 0.17 mM KCl, 0.33 mM CaCl2, 0.33 mM MgSO4, pH 7.4)). The following zebrafish lines were used in this study: AB/TL wild-type, *Tg*(*myl7*:*EGFP*)^twu3442^, and *wnt11^tx226^* mutant line^71^.

#### Cell culture

H9c2(2-1) (rat [Rattus norvegicus] heart myoblast) cells were obtained from ATCC^®^ (CRL-1446^™^), cultured in Dulbecco’s Modified Eagle Medium (DMEM) supplemented with 10% of Fetal Bovine Serum (FBS), 1% of Penicillin-Streptomycin and 1% of non-essential amino acids, and incubated at 37°C with 5% CO_2_. Neonatal rat ventricular myocytes (NRVMs) were isolated from 2 days old Wistar rat pups with digestion buffer (80µg/ml Liberase^TM^, 0.1% Trypsin, 20µg/ml DNase I, 10 µM CaCl2 in Hank’s Balanced Salt Solution), and maintained in DMEM-F12 medium with 10% foetal bovine serum and 1% penicillin/streptomycin for 24h hours, than cultured for next 48h with serum-free medium containing 80% DMEM-F12, 20% M199 with GlutaMAX Supplement and 1% Penicillin-Streptomycin.

C2C12 cells were a kind gift from the Birchmeier lab (MDC Berlin-Buch in the Helmholtz association, Germany). Cells were cultured in SILAC DMEM Medium (PAA Laboratories) at 37°C and 5% CO_2_ and complemented with 10% dFCS (Sigma), 1% Penicillin-Streptomycin (Gibco), 1x L-Glutamine (PAA M11-004), 1x L-arginine (Sigma) and 1x L-Lysine (Sigma) as described in^72^.

### METHODS DETAILS

#### siRNA-mediated Knockdown

Before transfection, NRVMs were switched to antibiotic-free culture medium. H9c2 cells were plated the evening before transfection, either on 12mm PDL-coated coverslips in a 24-well plate (5×10^4^ cells/ well), or 100mm petri dishes (2×10^6^/dish). All cells were transfected with ON-TARGETplus SMARTpool siRNA (Dharmacon) for 24h (H9c2), or 24-48h (NRVM) with a final concentration of 50nM per siRNA using DharmaFECT1 according to manufacturer’s instructions. To check the baseline response to siRNA treatment, a non-targeting “Scramble” siRNA was used as control.

#### qPCR

Total RNA was isolated from cultured H9c2 cells, NRMVs, and from 54 hpf zebrafish FACS-sorted cells using TRIzol according to the manufactur’s instructions. Isolated RNA was DNAse-treated with the DNase set (Qiagen), column purified using RNeasy Mini Kit (Qiagen), and reverse transcribed using First Strand cDNA Synthesis Kit (Thermo Fisher). 10-100 ng of cDNA was used for real-Relative mRNA levels were quantified by gene-specific TaqMan assays (see Table S4) using ViiA^TM^ 7 Real-Time PCR System and Quant Studio 7. Each reaction was performed in triplicates. Reactions with no template and no reverse transcriptase were included as controls. Threshold cycle (C_T_) values were normalized to those of relevant housekeeping genes, and fold change was quantified using ΔΔCt method ^73^. ΔC_T_ values matched by specific experiment were analysed with two-tail Wilcoxon signed rank test. To quantify gene knock-down efficacy of all targeting siRNA, paired ΔC_T_ values were analysed with one-tail Wilcoxon signed rank test. All results were plotted as log_2_ of fold change (2^-ΔΔCt).

#### Single-cell dissociation and FACS

54 hpf *Tg*(*myl7:EGFP*)^twu34^ embryos were dechorionated using Pronase, anesthetized using 0.016% tricaine (w/v) (pH∼7), and incubated in 0.25% Trypsin at 28.5°C. Single-cell dissociation was monitored every 20 minutes and was generally achieved within 2 hours. Efficient dissociation was facilitated by gentle pipetting. Cell suspension was filtered through a 70 μM strainer, and cells were thoroughly washed three times in cold PBS supplemented with Fetal Calf Serum in reducing concentration, and eventually resuspended in 1 ml of cold PBS. Single cells were then filtered in a glass tube with a cell strainer cap and placed on ice until counting. Samples were sorted using a BD FACS Aria1 flow cytometer, operating with BD LSRFortessa analyzer with BD FACSDiva software. GFP-expressing cells were sorted using a 488nm laser and a 530/30 BP filter.

#### Drug treatment

H9c2 cells were plated 48 hours prior to drug experiments and incubated with the following: 10 μM Foskolin (FSK), 10 μM Protein kinase inhibitor (PKI), 10 μM H-89, 10 μM Okadaic acid (OA), 10 μM Calpeptin (CLP), 100 μM L314E for 1 hour or with 4 μM isopropanol (ISO) for 3 minutes, and with corresponding concentration of DMSO as a control before harvesting or fixation.

#### Ratriometric FRET

Relative PKA activity in cells were estimated using AKAR4-NES, a cytosol-targeted FRET-based PKA biosensor. H9c2 cells were plated on 8-well plates (µ-Plate, ibidi) at a density of 2×10^4^ cells/well, and transfected with 300ng of the sensor using Lipofectamine^TM^ 2000 per manufacturer’s instructions. Cells were imaged 48h post-transfection in FluoroBrite^TM^ DMEM supplemented with 10% FBS on an Olympus IX81 microscope, equipped with a UV Apochromat 20X/0.75 DIC objective. Dual emission imaging was performed using a 430/25 excitation filter, a zt442RDC dichroic mirror, and a 483/32 (for CFP) or a 542/27 (for FRET acceptor channel) emission filter as appropriate. All images were automatically acquired with an ImagEM CCD 9100-13 camera (Hamamatsu) at 16-bit depth on a motorized stage (Märhäuser Corvus-2) and automated ZDC z-Focus sytem. Images were processed using Fiji to calculate FRET ratios. Average background images were calculated for each channel from untrasfected control cells, and were substracted from raw image data to correct for background. Region of interests (ROI) were defined from thresholded images of each channel, and combined using the conjunction operation “AND” to generate the final ROI composite. FRET ratios were calculated within the final ROI, and normalized to the maximum value for each frame and to the average of untreated control. Mean differences between two groups were analyzed with a two-tail unpaired t-test with Welch’s correction.

#### Immunofluorescence staining and confocal microscopy

H9c2 cells were plated on poly-D-lysine coated (0.1mg/mL, Sigma) coverslips 48 hours prior to the experiments, and fixed for 20 minutes in 4% PFA supplemented with 4% Sucrose. Cells were permeablized with 0.2% Triton X-100. These following antibodies were diluted in 1% Gelatine at the specified dilutions: anti-II-III loop (Millipore) 1:100, Alexa Fluor® 633 phalloidin (Thermo Fisher Scientific) 1:4, anti-pCt (Abnova) 1:500, anti-nucleolin (Abcam) 1:500. Z-stacks of images were obtained on a Leica TCS SP8 confocal microscope using the HC PL APO 63x/1.40 OIL CS 2 objective (NA =1.4).

Fluorescence intensity from maximal z-projections was quantified using ImageJ/Fiji. To assess changes in pCt accumulation within nucleoli, we made a mask using thresholded signals from Nucleolin, and mean fluorescence intensity values from pCt signals were then measured in individual objects larger than four pixels. All values were then corrected for background signals before normalized to the inverse of average intensities from control untreated cells 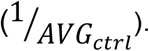 Mean differences between two groups were analyzed with a two-tail unpaired t-test with Welch’s correction for unequal variance as appropriate. For analysis of difference among ≥3 groups, we used a standard One-Way ANOVA with Tukey’s correction for multiple comparison of mean values from treatment groups with the mean value from control. Additionally, frequency distribution of all normalized values were visualized as density curves and/or histogram with equal bandwidth/binwidth of 0.2.

#### Immunoblotting

H9c2 cell lysates were prepared using RIPA buffer (20 mM Tris-HCl, 150 mM NaCl, 0.1% Triton X-100, 0.1% SDS, 0.5% Sodium-deoxycholate, PhosSTOP, EDTA-free Protease Inhibitor Cocktail). Lysates were sonificated and centrifuged at 5000x*g* at 4°C for 5 minutes. Protein concentrations were quantified by Lowry’s estimation method. After addition of Laemmli buffer (50 mM Tris-HCl, 0.1% SDS, 9% Glycerol, 1% β-mercaptoethanol, 0,02% Bromphenolblue, 25 mM DTT) lysates were cooked for 10 minutes and centrifuged again at 5000x*g* and 4°C for 5 minutes. Total protein was separated on 4-15% SDS-polyacrylamide gel (BioRad), and transferred to a Nitrocellulose Blotting membrane (GE Healthcare). Membranes were blocked for 1 hour in 5% milk/TBS-T. The following primary antibodies’ were diluted in blocking buffer and used in the specified dilutions: anti-AKAP2 (Bioscience) 1:1000, anti-β-actin (Sigma) 1:3000, and anti-pCt (custom-made, Cambridge Research Biochemicals) 1:1000. Membranes were incubated with primary antibodies with gentle agitation overnight at 4°C, followed by incubation with appropriate HRP-coupled secondary antibodies. Chemiluminescent detection was performed using Pierce ECL Western Blotting Substrate (Thermo Scientific). All western blots were quantified with ImageJ/Fiji.

#### Plasmid construction

cDNA encoding FLAG-CT-mVenus of rat *CACNA1C* and rat *AKAP2* was purchased from Thermofisher Scientific. FLAG/HA-tagged AKAP2 was cloned using Gateway technology (Invitrogen) by PCR amplification of the *AKAP2* cDNA using attB-flanked BP primers (see Table S2). attB-flanked PCR product was cloned first into a pDONR™221 vector using BP Clonase™ II Enzyme mix to generate an entry vector, and subsequently transferring into a destination vector pFRT_TO_DESTFLAGHA to create expression clone using LR Clonase™ II Enzyme mix.

#### Immunoprecipitation

H9c2 cells were transfected with a concentration of 200 ng/ml plasmid DNA using Lipofectamine™ 2000 according to manufacturer’s instructions. After 24h, cells were solubilized in mild lysis buffer (2mM EDTA, 2mM EGTA, Triton X-100 in PBS) containing PhosSTOP and EDTA-free Protease Inhibitor Cocktail. Lysates were passed through a syringe of 0.4mm diameter, centrifuged at 5000xg at 4°C for 5 minutes and quantified by Bradford’s protein estimation method (Anal. Biochem. 72, 248-254 (1976)). Lysates were incubated over night at 4°C either with Anti-FLAG M2 Magnetic Beads or with Dynabeads Protein A, which was beforehand incubated with AKAP2 or GFP antibody. Beads were washed four times with mild lysis buffer using magnetic separator. Immunoprecipitated proteins were eluted with 0.1M glycine (pH 2.5), neutralized with 1M Tris (pH 10.6) and denatured in 1x Laemmli sample buffer (10 % glycerol; 2 % SDS; 0.01 % bromophenol blue; 75 mM Tris-HCl; 50 mM DTT; pH 6.8) for 7 min at 95°C. Eluted proteins were resolved on 10% SDS-PAGE gels and western blotted using standard procedure. The following primary antibodies were used: anti-AKAP2 (Bioscience) 1:500, anti-FLAG (Sigma) 1:1000 and anti-pCt (custom-made, Cambridge Research Biochemicals) 1:1000, anti-GFP (GeneTex) 1:4000. Chemiluminescent detection was performed using Immobilon HRP substrate (Merck #WBKLS0500), and western blots were visualized using Odyssey Fc Imaging System (LI-COR Biosciences).

#### C2C12 cell sample preparation for Mass Spectrometry

For SILAC experiments the medium was alternatively supplemented with Arg6 and Lys4 (medium) or Arg10 and Lys8 (heavy). For the differentiation, cells were transferred to DMEM containing 2% dialyzed horse serum (Gibco), 1% Penicillin-Streptomycin (Gibco), 4mM Glut, L-Arg and L-Lys at 90% confluency. The differentiation medium was exchanged every second day. Differentiation was observed to start after three days and was finished at day 5. Cells were grown in light, medium and heavy conditions, and were harvested at different time points after start of differentiation (0h light, 10h light, 30h medium, 100h heavy, 150h heavy). Cells were washed with PBS, spun down and mixed to obtain two SILAC sample mixtures (sample1: 0h light, 30h medium and 100h heavy; sample2: 10h light, 30h medium and 150h heavy).

Cells were lysed with NuPage LDS Sample Buffer (Invitrogen) and 150µg protein per sample were separated by SDS-Page on a 4-12% SDS-polyacrylamid gel (Invitrogen). Proteins were fixed and stained using a colloidal blue staining kit (Invitrogen). Gel pieces were washed sequentially with ABC (50 mM AmmoniumBiCarbonate), ABC/EtOH and EtOH for 10 min each at room temperature. Gel pieces were dried in a speed vac for 5 min at 45°C, got rehydrated with 10 mM DTT in 50 mM ABC and incubated for 60 min at 56°C. Gel pieces were incubated for 45 min in 55 mM iodacetamide and 50 mM ABC buffer in the dark, then they were washed once with ABC at room temperature, and dehydrated twice. Remaining EtOH was removed by vacuum centrifugation, trypsin solution added (0.5µg/µl in 50mM ABC) and left shaking overnight at 37°C. The liquid was transferred to a fresh tube, peptides were extracted from gel pieces by adding extraction buffer (3% TFA, 30% acetonitrile) and liquid was transferred to the corresponding fresh tubes. Pure acetonitrile (LC-MS grade, Merck) was added for 10 min, supernatant additionally added to the corresponding tubes and samples vacuum dried to 10-20 % of the original volume to remove acetonitrile. Fractions were desalted on stage tips as described ^74^. Vacuum dried peptide fractions were resuspended (5 % Acetonitrile and 0.1 % formic acid) and analyzed by mass spectrometry.

#### Mass Spectrometry analysis

Peptides were separated on a High Performance Liquid Chromatography System (Thermo Scientific) using a 15 cm column (75 µm inner diameter) packed in house with ReporSil-Pur C18-AQ material (Dr. Maisch, GmbH). The applied reverse phase gradient ran from 5 to 60 % Acetonitrile within 3 h. Peptides were ionized with an electrospray ionization source (Thermo Scientific) and analyzed on a Q Exactive mass spectrometer (Thermo Scientific). The mass spectrometer was running in data-dependent mode to select the 10 most intense ions for fragmentation from the corresponding MS full scan. Other parameters were: 70,000 ms1 resolution; 3,000,000 ions target value for ms1; maximum injection time of 20 ms; 17,500 ms2 resolution; 60 ms maximum ion collection time; 1,000,000 ions target value for ms2.

#### Microinjection

Morpholinos (MO) were dissolved in RNase/Dnase-free water to make 1mM stock solutions, and used at the specfied dilutions: *akap2* MO 1:1 (e1i1, splice MO) or 1:2 (ATG MO), *wnt11* MO 1:6. Diluted MO solutions were incubated at 65°C for 10 minutes before injection into the yolk of 1-4 cell stage embryos. As negative controls, we also used standard control MO and mismatch MO, both at a 1:1 dilution. All MO used were obtained from GeneTools.

Capped *akap2* mRNA or *akap2ΔPKA* mRNA (75 pg/μl each) was injected into 1-cell stage embryos. sgRNA-Cas9 injection mix was prepared as descibed^75^: *akap2* sgRNA (200ng/μl) was mixed with purified Cas9 protein(600ng/µl), and supplemented with 0.2M KCl. All solutions were diluted in RNAse-and DNAse-free water (Sigma). Injection volume is 1nl per embryo for all experiments.

#### mRNA construction

Zebrafish *akap2* (ENSDARG00000069608) mRNA was constructed by PCR amplification of cDNA using *Danio rerio* akap2 infusion primers (see Supplementary Table 2), which were designed with http://bioinfo.clontech.com/infusion/convertPcrPrimersInit.do. The cDNA fragment was cloned into pBluescript KS (-) vector with HindIII at the 5’-end and BamHI restriction enzyme site at the 3’-end using the In-Fusion® HD Cloning System. *In vitro* transcription was performed with mMessage mMachine T3 Transcription Kit using BamHI-linearized DNA template at 37°C. To increase transcript stability and translation efficiency, we performed polyadenylation of the mRNA using Poly(A) Tailing Kit. *Akap2* cDNA lacking the PKA binding site (akap2ΔPKA) was generated by PCR-mediated deletion of plasmid DNA using the full-length cDNA construct described above as template. The primer pair was designed corresponding to 30 nucleotides (nt) up and downstream from the PKA binding sequence of 75nt. The construct was amplified by *Pfu* DNA polymerase according to manufacturer’s instructions in a 50µl PCR reaction containing 2mM Mg^2+^. Template plasmid was removed by DpnI digestion, followed by PCR clean-up and ligation by T4 ligase. The vector was then cloned into chemically competent DH5α *E.coli*. After confirming successful PKA binding site deletion from isolated plasmid DNA by sequencing, the plasmid was linearized by BamHI before RNA synthesis with T7 MAXIscript® Kit. akap2ΔPKA mRNA was again polyadenylated using Poly(A) Tailing Kit, and purified with RNeasy Mini Kit before injection.

#### CRISPR/Cas9-mediated Mosaic Mutagenesis

For targeted mutagenesis of zebrafish embryos using the CRISPR/Cas9 system, gRNA targeting exon 2 of *Akap2* was designed with <u>chopchop.cbu.uib.no</u>. We synthesized sgRNA by in vitro transcription with MEGAscript^TM^ T7 Transcription Kit using sgRNA primers (see Table S2). Genomic DNA was isolated from wild-type, Cas9 or sgRNA-Cas9 injected 54 hpf single embryos and amplified with flanking primers (see table S2) designed to amplify the sgRNA target site, and subsequently cloned into pGEM®-T Easy Vector. To analyze specific allele variations, we used CrispRVariantsLite as described^76^.

#### Zebrafish embryos imaging

54 hpf embryos were either anesthetized using 0.016% tricaine (w/v) (pH∼7), or fixed in PEM solution (100 mM PIPES, 2 mM MgSO_4_, 1mM EGTA, pH 7) with 4% Formaldehyde and 0.1% Triton X-100 for 2 h at RT. Embryos were mounted in 2% methylcellulose for imaging using a Leica M165 Fluorescence Stereo Microscope equipped with a Leica CCD camera. Images were processed using ImageJ/Fiji and Adobe Photoshop CS6.

#### Whole-mount In situ Hybridization (ISH)

Whole-mount ISH was carried out as described previously^77^. Briefly, both *akap2* anti-sense and sense DIG-labeled probes were synthesized from cDNA template by T7 polymerase. Dechorinated and PTU-treated embryos were fixed in 4%PFA. Permeabilized embryos (10ug/ml Proteinase K, 20min) were labeled with Alkaline phosphatse anti-DIG antibody diluted in blocking buffer (2mg/ml BSA in PBT) at a 1:5000 dilution overnight at 4°C. Labeling reactions in NBT/BCIP solution occured in the dark for 3-5 hours until the desired intensity of signals was achieved. Embryos mounting and imaging were done as described.

#### RT-PCR

Total RNA was isolated from cultured H9c2 cells, and from 54 hpf zebrafish embryos using the TRIzol reagent according to the manufactur’s instruction. RNA was reverse transcribed using First Strand cDNA Synthesis Kit and amplified using Phusion polymerase (NEB) with primers for the relevant genes (see Table S2). The PCR products were visualized on 3% agarose gel stained with Redsafe.

#### Heart Looping Measurement

Heart angle was defined as the angle enclosed by the midsagittal line and the atrioventricular canal as described^78^. For the measurements, fixed 54 hpf *Tg*(*myl7*:*EGFP*)^twu34^ embryos were imaged as described. Images were analyzed using ImageJ/Fiji.

#### Confocal [Ca2+]_i_ imaging in isolated NRVM

NRVMs were loaded with Ca^2+^ indicator Fluo-4 AM as previously described^79^, transferred to the stage of a confocal microscope (Zeiss LSM800, excitation 488 nm, emission >515 nm) and superfused with Tyrode’s solution (in mM: NaCl 130, KCl 4, CaCl2 1.8, MgCl2 1, D-glucose 10, HEPES 10; pH 7.4 with NaOH). NRVMs were stimulated in an electrical field (1 Hz) and cytosolic Ca^2+^ transients were recorded along a line repetitively scanned at the equatorial plane along the long axis of the cell (pixel size: 0.07-0.13, 0.8 ms/line, respectively). The global Ca^2+^ transient was calculated from the average intensity of the line within the cell recorded over time. Local Ca^2+^ release was quantified in 1 µm intervals along the line. Local Ca^2+^ increase was defined as „late” if local [Ca^2+^] reached half-maximum of the global Ca^2+^ transient (F50) in >6 ms, or „early” if local time to F50 was ≤6 ms, indicating close proximity to an active couplon/Ca^2+^ release unit^80^. For early sites, i.e. with early Ca^2+^ release in at least 1 out of 10 consecutive Ca^2+^ transients at steady state, open probability of the couplon (P_o_) was calculated by dividing the number of transients with early release by the number of total consecutive transients recorded.

#### High-speed Ratriometric Calcium Imaging in Isolated Embryonic Zebrafish Hearts

Recording of the intracellular Ca^2+^ transients in developing zebrafish heart was perofrmed as descirbed^32^. Hearts were isolated from 54 hpf zebrafish embryos in normal Tyrode’s solution (NTS, see below) supplemented with 20 mg/ml BSA. For ratiometric calcium transient recordings, hearts were loaded for 15 min with 50mM of the calcium-sensitive dye Fura-2AM (Thermo Fisher Scientific) and subsequently washed in dye-free NTS. Hearts were then incubated in NTS at room temperature for 30–45 min to allow complete intracellular hydrolysis of the esterified dye. A high-speed monochromator (Optoscan, Cairn) was used to switch the excitation wavelength rapidly between 340 nm and 380 nm with a bandwidth of 20 nm and at a frequency of 500 s^-1^. The excitation light was reflected by a 400-nm cut-off dichroic mirror and fluorescence emission was collected by the camera through a 510/80-nm emission filter. High-speed CCD camera (RedShirtImaging) and equipped with NeuroPlex software was used for recordings. Images were analyzed with MatLab (MathWork) using customized software^32^.

#### Optical mapping of action potential propagation

Optical mapping and signal processing was performed as previously described^32^. Hearts were isolated from 54 hpf zebrafish embryos in normal Tyrode’s solution (NTS contains 136 mM NaCl, 5,4 mM KCl, 1mM MgCl_2_x6H_2_O, 5mM D-(+)-Glucose, 10mM HEPES, 0.3 mM Na_2_HPO_4_x2 H_2_O, 1. mM CaCl_2_x2 H_2_O, pH 7.4) supplemented with 20 mg/ml BSA. To record membrane potential changes, hearts were stained for 12 minus FluoVolt probe (Thermo Fisher Scientific), and washed with NTS-BSA. Individual hearts were transferred into perfusion bath (Warner Instruments), which contained NST supplemented with 100 μM Cytochalasin D to inhibit contraction. For the imaging high-speed CCD camera (RedShirtImaging) was used equipped with 488 nm LED lamp and NeuroPlex software. Images were analyzed with MatLab (MathWork) using customized software^32^.

### QUANTIFICATION AND STATISTICAL ANALYSIS

Data analysis and visualization were done using GraphPad Prism 7 and R. For all statistical tests, significance level (alpha) was set at 0.05. All p-values >0.0001 are rounded to the third decimal. Specific p-values <0.05 are included in the figures, and p-values>0.05 are in corresponding figure legends. Equal variance of normally distributed data was checked with an F-test. Differences between groups were analyzed as indicated in corresponding method sections. Experiment-specific statistical details can be found within figures and figure legends.

#### Analysis of mass spectrometry data

Raw files were analyzed with MaxQuant 1.6.0.1^81^. A mouse specific database (Uniprot 2018-02, canonical and isoforms) was used for the peptide search. Multiplicity was set to 3, Arg6 and Lys4 defined as medium and Arg10 and Lys8 as heavy. Trypsin was set as the protease including the option to cut after proline. Deamidation at N and Q, N-terminal acetylation and oxidation at methionine were set as variable modifications. Carbamidomethylation of cysteines was set as a fixed modification. Peptide and protein FDR were set to 0.01. The proteinGroups.txt file was used for further processing of the data in R. The table was prefiltered to exclude proteins with a peptide count less than two and also excluding potential contaminants. The timeseries was extracted from normalized ratios by using the 30h timepoint as a reference. Therefore, the 30h timepoint for every given protein is set to 0 while changes in other timepoints are expressed as relative values to 0. The timepoints at 0h and 100h were considered as representative timepoints for myoblasts and myotubes respectively, and log2-fold changes between both timepoints were plotted. iBAQ values as calculated by MaxQuant^72^ were used to approximate overall relative protein abundances.

#### Data availability

The data that support the findings in this study are available within this article and its Supplementary Information files, and from the corresponding author upon reasonable request.

---

**Supplementary Figure 1.**
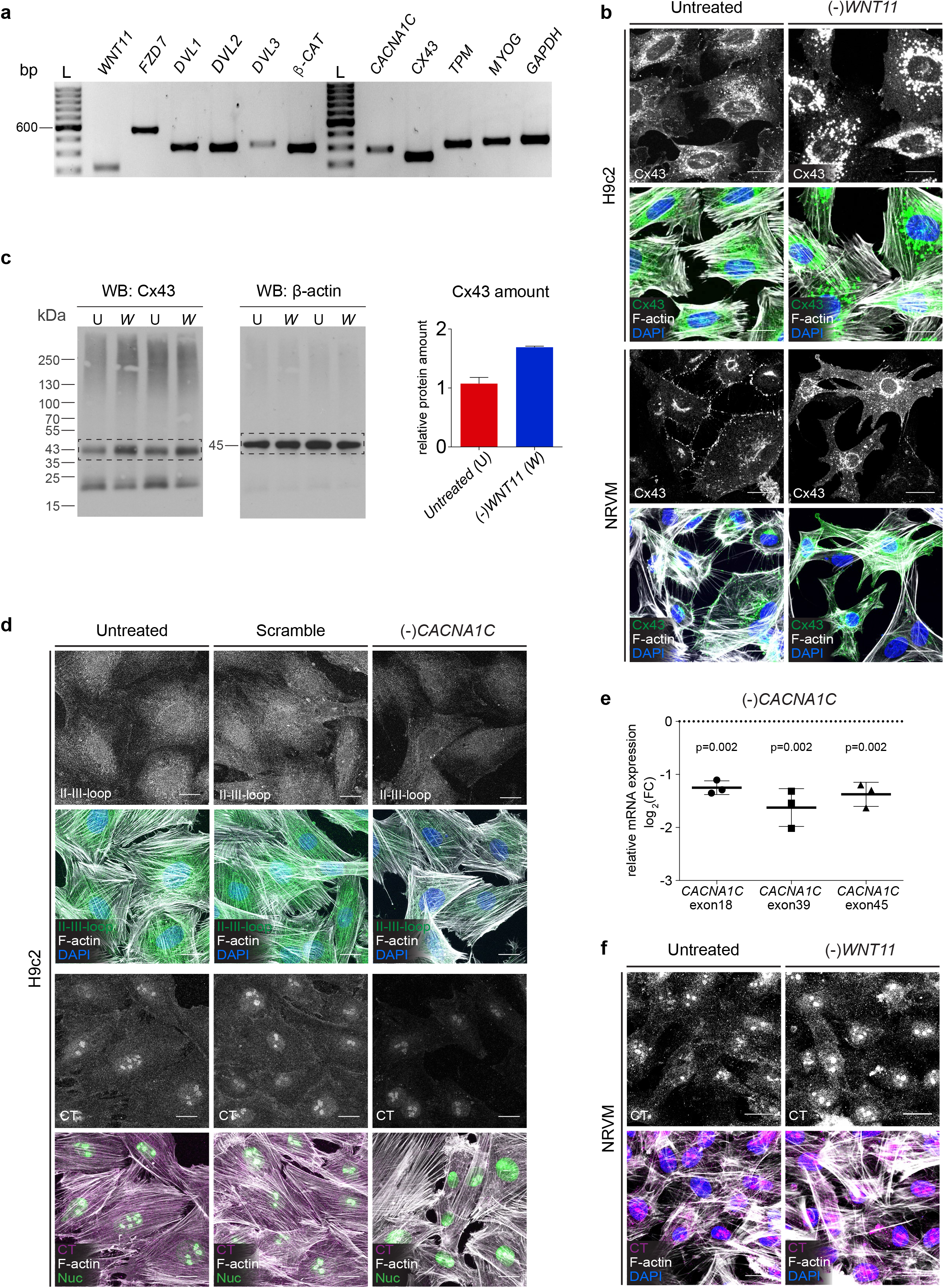
**a** RT-PCR of cDNA of the indicated gene from H9c2 cells. **b** Representative maximum intensity z-projection of Cx-43 signals in untreated, and *WNT11* siRNA-transfected H9c2 or NRVM cells alone (gray scale), and merged (green) with F-actin signals (gray). Scale bar, 20 μm. **c** Western blot (WB) detection of Connexin 43 (Cx43) in lysates of untreated control (U) and *WNT11* siRNA transfected (*W*) H9c2 cells using anti-Cx43 antibody. On the right, column graph represent the semi-quantified Cx43 amount normalized to β-actin. (Mean ± SD; N = 2). **d** Representative maximum intensity Z-projections of II-III-loop signals in untreated, scramble and *CACNA1C* siRNA transfected H9c2 cells alone (gray scale), and merged (green) with F-actin signals (gray); and of CT signals in untreated, scramble and *CACNA1C* siRNA transfected H9c2 cells alone (gray scale), and merged (magenta) with nucleolin (Nuc) (green), and F-actin (gray) signals. Scale bar, 20 μm. **e** Quantification of relative mRNA expression of *CACNA1C*, using Taqman probes targeting three different exons in scramble and *CACNA1C* siRNA-transfected H9c2 cells. Data plotted as log_2_ of fold change ± SD of N = 3 experiments. One-tailed Wilcoxon rank sum test. **f** Representative maximum intensity z-projections of CT signals in untreated, and *WNT11* siRNA-transfected NRVM cells (gray scale), and merged (magenta) with F-actin signals (gray). Scale bar, 20 μm.

**Supplementary Figure 2.**
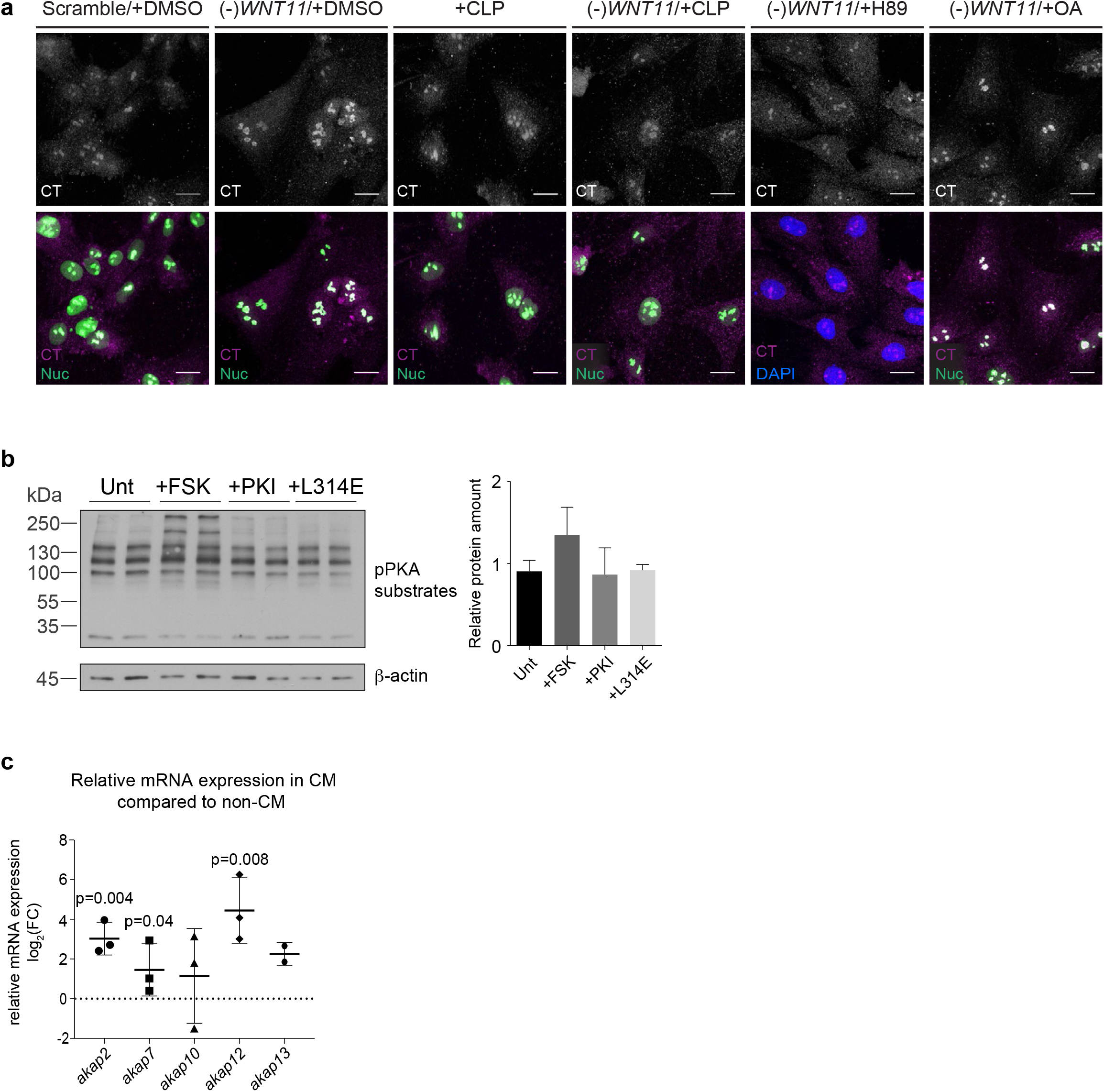
**a** Representative maximum intensity z-projection of CT signals in scramble, or *WNT11* siRNA­ transfected H9c2 cells treated with 1% DMSO, Calpeptin (CLP, S0 tg/ml), H89 (10µM), or with Okadaic acid (OA, 10µM) (gray scale), and merged (magenta) with nucleolin (Nuc) (green). Scale bar, 20 µm. **b** Western blot (WB) detection of phosphorylated PKA substrates level in untreated control (Unt), and of FSK, PKI or L314E-treated H9c2 cells. Bar graph (right) represents semi-quanittative phosphorylated PKA substrates amount, normalized to 13-actin. (N =2 experiments). c Quantification of relative mRNA expression of preselected *akap* genes in sorted embryonic zebrafish cardiomyocytes (CM) compared to non-CM. Data plotted as 109 2 of fold change (FC) ± SD of N = 3 experiments. Wilcoxon rank sum test; *Pakap1o=0.078, Pakap13=0.06 3*.

**Supplementary Figure 3.**
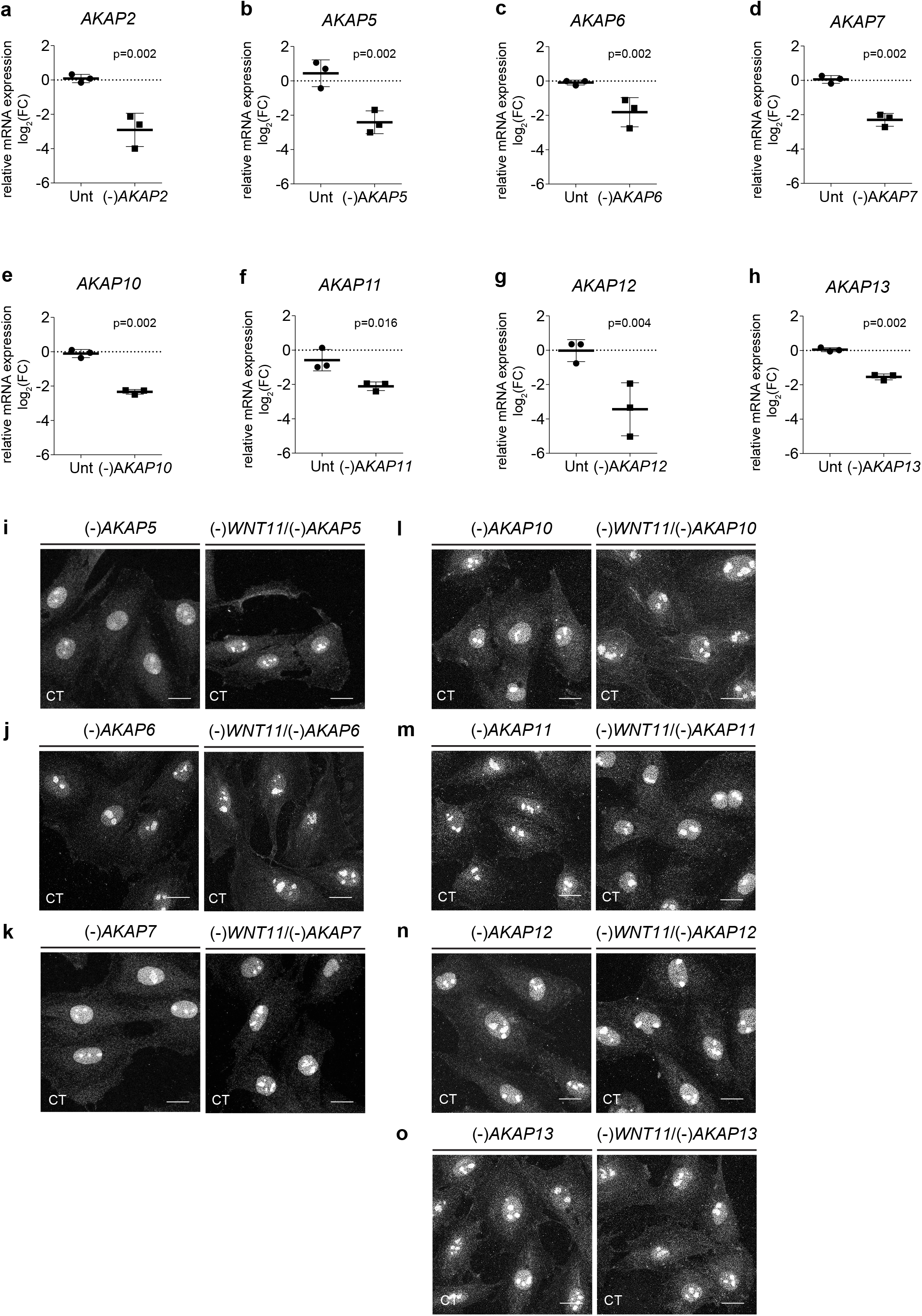
**a-h** Quantification of relative mRNA expression of untreated, scramble, *AKAP* siRNA-treated H9c2 cells showing siRNA efficacy. Data plotted as log_2_ of fold change (FC) ± SD of N = 3 experiments. Wilcoxon rank sum test; *AKAP2*: *P*_Unt_=0.426, *AKAP5*: *P*_Unt_*=*0.074, *AKAP6*: *P*_Unt_*=*0.36, *AKAP7*: *P*_Unt_*=*0.301, *AKAP10*: *P*_Unt_*=*0.159, *AKAP11*: *P*_Unt_*=*0.219, *AKAP12*: *P*_Unt_*=*0.945, *AKAP13*: *P*_Unt_*=*0.426. **i-o** Representative maximum intensity z-projection of CT signals (gray scale) in *AKAP*, or AKAP/*WNT11* siRNA-transfected H9c2 cells. Scale bar, 20 μm.

**Supplementary Figure 4.**
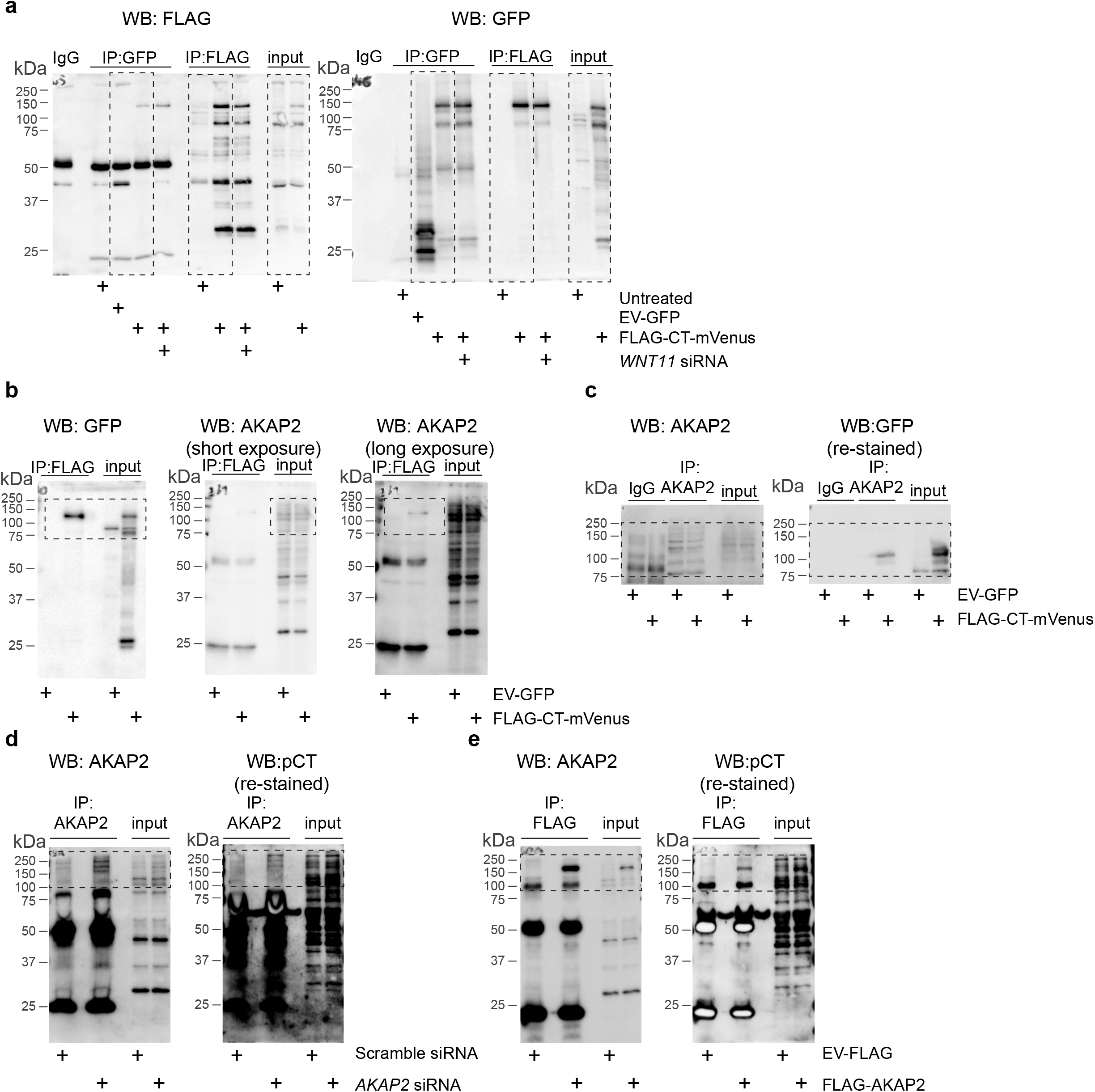
**a** FLAG or GFP detection by WB of the FLAG-and mVenus-tagged CT construct from lysate of H9c2 cells either transfected with the construct or with empty vector GFP (EV-GFP) control, before and after FLAG or GFP IP. (N = ≥ 10 experiments). **b** GFP or AKAP2 Co-IP detection by WB of CT construct in of lysates of H9c2 cells, transfected either with the CT construct or with empty vector GFP (EV-GFP) control, showing input and FLAG-IP. **c** Endogenous AKAP2 (left) or GFP (right) Co-IP detection by WB lysates of H9c2 cells, transfected either with the CT construct or with empty vector GFP (EV-GFP) control. **d** Endogenous AKAP2 and CT Co-IP detection is lysates from Scramble or *AKAP2* siRNA transfected H9c2 cells, using anti-AKAP2 or anti-pCT antibody as noted. **e** AKAP2 and CT Co-IP detection in lysates from H9c2 cells transfected with FLAG/HA-tagged AKAP2 construct or EV-FLAG, showing input and FLAG-IP.

**Supplementary Figure 5.**
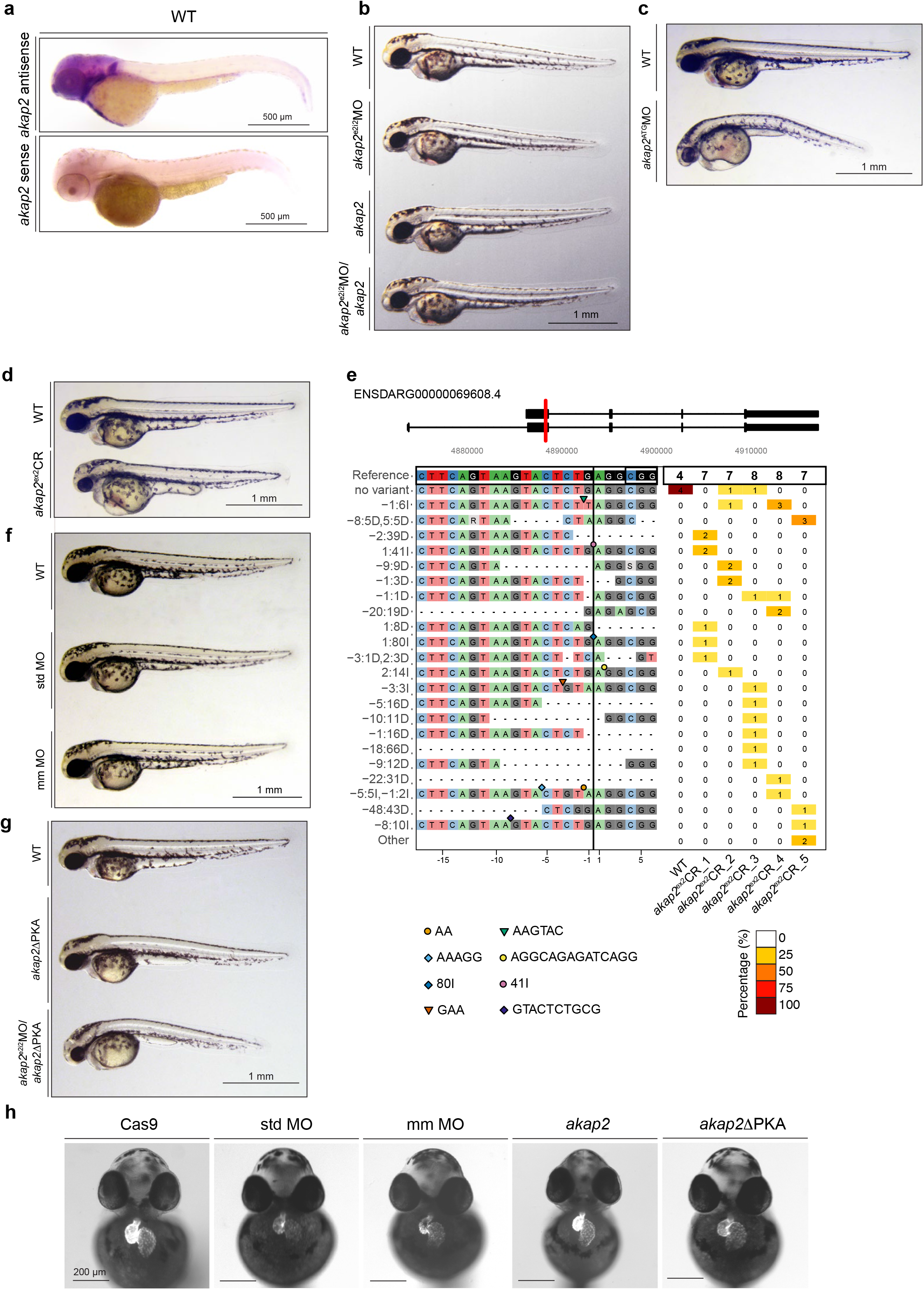
**a** Bright field images (lateral view) of *in situ* hybridization detection of endogenous *akap2* mRNA using antisense (upper panel) and sense (lower panel) *akap2* probe in wild-type (WT) zebrafish embryos at 48 hpf. **b** Bright field images (lateral view) of wild-type (WT), *akap2* splicing site targeting morpholino (*akap2*^e2i2^MO), *akap2* mRNA (*akap2*), and *akap2* morpoholino/mRNA (*akap2*^e2i2^MO/*akap2*) double-injected zebrafish embryos at 54 hpf. **c** Bright field images (lateral view) of wild-type (WT) and *akap2* start site targeting morpholino (*akap2*^ATG^MO) injected zebrafish embryos at 54 hpf. **d** Bright field images (lateral view) of wild-type (WT) and *akap2* sgRNA/Cas9 (*akap2*^ex2^CR) injected zebrafish embryos at 54 hpf. **e** Panel plot showing allele variations in CRISPR/Cas9 (*akap2*^ex2^CR) injected embryos compared to WT *akap2* allele (ENSDARG00000069608). **f** Bright field images (lateral view) of wild-type (WT), standard (std MO), and mismatch (mm MO) control morpholino injected zebrafish embryos at 54hpf. **g** Bright field images (lateral view) of wild-type (WT), *akap2*ΔPKA mRNA, *akap2*ΔPKA mRNA and *akap2* double (*akap2*^e2i2^MO/*akap2*ΔPKA) injected zebrafish embryos at 54 hpf. **h** Bright field images overlayed with fluorescent images (frontal view) of Cas9, standard (std MO), mismatch (mm MO), *akap2* mRNA, *akap2*ΔPKA mRNA injected *myl7*:*EGFP* transgenic zebrafish embryos at 54hpf. Scale bar=200 μm.

**Supplementary Figure 6.**
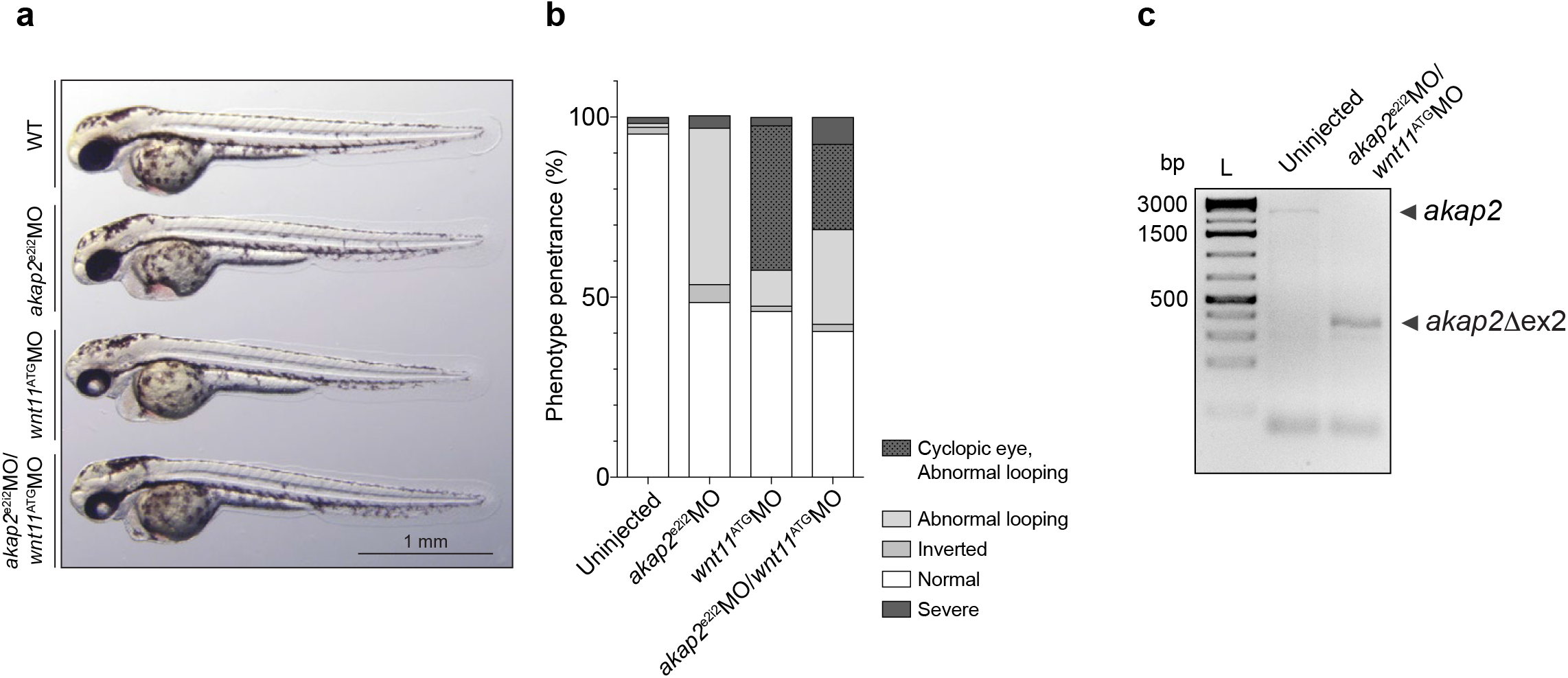
**a** Bright field images (lateral view) of wild-type (WT), *akap2* splicing site targeting morpholino (*akap*2^e2i2^MO), *wnt11* morpholino (*wnt11*^ATG^MO), and double-injected (*akap*2^e2i2^MO / *wnt11*^ATG^MO) zebrafish embryos at 54 hpf. **b** Quantification of single-and double-injected (*akap*2^e2i2^MO / *wnt11*^ATG^MO) phenotypes based on heart looping abnormalities and cyclopic eye. (N ≥ 3 experiments; n ≥ 300). **c** RT-PCR of uninjected and double-injected (*akap*2^e2i2^MO / *wnt11*^ATG^MO) zebrafish embryos at 54hpf. The size of the akap2 fragment is 2434 bp, while in *wnt*11/*akap2* morphants it is only 300bp due to exclusion of the exon 2 (*akap2*Δex2)

## References

1. Bers, D. M. Cardiac excitation-contraction coupling. Nature 415, 198–205 (2002).

2. Carafoli, E., Santella, L., Branca, D. & Brini, M. Generation, control, and processing of cellular calcium signals. Crit. Rev. Biochem. Mol. Biol. 36, 107–260 (2001).

3. Catterall, W. A. Structure and regulation of voltage-gated Ca2+ channels. Annu. Rev. Cell Dev. Biol. 16, 521–555 (2000).

4. Zühlke, R. D., Pitt, G. S., Deisseroth, K., Tsien, R. W. & Reuter, H. Calmodulin supports both inactivation and facilitation of L-type calcium channels. Nature 399, 159–162 (1999).

5. Catterall, W. A. Voltage-Gated Calcium Channels. Cold Spring Harb Perspect Biol 3:a003947 (2011).

6. van der Heyden, M. A. G., Wijnhoven, T. J. M. & Opthof, T. Molecular aspects of adrenergic modulation of cardiac L-type Ca2+ channels. Cardiovasc. Res. 65, 28–39 (2005).

7. Weiss, S., Oz, S., Benmocha, A. & Dascal, N. Regulation of cardiac L-type Ca²⁺ channel CaV1.2 via the β-adrenergic-cAMP-protein kinase A pathway: old dogmas, advances, and new uncertainties. Circulation research 113, 617–631 (2013).

8. De Jongh, K. S. et al. Specific Phosphorylation of a Site in the Full-Length Form of the α1 Subunit of the Cardiac L-Type Calcium Channel by Adenosine 3‘,5‘-Cyclic Monophosphate-Dependent Protein Kinase †. Biochemistry 35, 10392–10402 (1996).

9. Gerhardstein, B. L. et al. Proteolytic processing of the C terminus of the alpha(1C) subunit of L-type calcium channels and the role of a proline-rich domain in membrane tethering of proteolytic fragments. J. Biol. Chem. 275, 8556–8563 (2000).

10. Gao, T. et al. C-terminal fragments of the alpha 1C (CaV1.2) subunit associate with and regulate L-type calcium channels containing C-terminal-truncated alpha 1C subunits. J. Biol. Chem. 276, 21089–21097 (2001).

11. Hulme, J. T., Yarov-Yarovoy, V., Lin, T. W.-C., Scheuer, T. & Catterall, W. A. Autoinhibitory control of the CaV1.2 channel by its proteolytically processed distal C-terminal domain. J. Physiol. (Lond.) 576, 87–102 (2006).

12. Fuller, M. D., Emrick, M. A., Sadilek, M., Scheuer, T. & Catterall, W. A. Molecular mechanism of calcium channel regulation in the fight-or-flight response. Sci Signal 3, ra70 (2010).

13. Edwards, A. S. & Scott, J. D. A-kinase anchoring proteins: protein kinase A and beyond. Curr. Opin. Cell Biol. 12, 217–221 (2000).

14. McConnachie, G., Langeberg, L. K. & Scott, J. D. AKAP signaling complexes: getting to the heart of the matter. Trends Mol Med 12, 317–323 (2006).

15. Dema, A., Perets, E., Schulz, M. S., Deák, V. A. & Klussmann, E. Pharmacological targeting of AKAP-directed compartmentalized cAMP signalling. Cell. Signal. 27, 2474–2487 (2015).

16. Hulme, J. T., Westenbroek, R. E., Scheuer, T. & Catterall, W. A. Phosphorylation of serine 1928 in the distal C-terminal domain of cardiac CaV1.2 channels during β1-adrenergic regulation. Proceedings of the National Academy of Sciences 103, 16574–16579 (2006).

17. Jones, B. W. et al. Cardiomyocytes from AKAP7 knockout mice respond normally to adrenergic stimulation. Proceedings of the National Academy of Sciences 109, 17099–17104 (2012).

18. Fuller, M. D., Fu, Y., Scheuer, T. & Catterall, W. A. Differential regulation of CaV1.2 channels by cAMP-dependent protein kinase bound to A-kinase anchoring proteins 15 and 79/150. J. Gen. Physiol. 143, 315–324 (2014).

19. Nichols, C. B. et al. Sympathetic stimulation of adult cardiomyocytes requires association of AKAP5 with a subpopulation of L-type calcium channels. Circulation research 107, 747–756 (2010).

20. Oliveria, S. F., Dell’Acqua, M. L. & Sather, W. A. AKAP79/150 anchoring of calcineurin controls neuronal L-type Ca2+ channel activity and nuclear signaling. Neuron 55, 261–275 (2007).

21. Hulme, J. T., Lin, T. W.-C., Westenbroek, R. E., Scheuer, T. & Catterall, W. A. Beta-adrenergic regulation requires direct anchoring of PKA to cardiac CaV1.2 channels via a leucine zipper interaction with A kinase-anchoring protein 15. Proc. Natl. Acad. Sci. U.S.A. 100, 13093–13098 (2003).

22. Oz, S. et al. Protein kinase A regulates C-terminally truncated CaV 1.2 in Xenopus oocytes: roles of N-and C-termini of the α1C subunit. J. Physiol. (Lond.) 595, 3181–3202 (2017).

23. Katchman, A. et al. Proteolytic cleavage and PKA phosphorylation of α1C subunit are not required for adrenergic regulation of CaV1.2 in the heart. Proceedings of the National Academy of Sciences 114, 9194–9199 (2017).

24. Schulte, G. International Union of Basic and Clinical Pharmacology. LXXX. The class Frizzled receptors. Pharmacol. Rev. 62, 632–667 (2010).

25. Wright, S. C. et al. A conserved molecular switch in Class F receptors regulates receptor activation and pathway selection. Nat Commun 10, 667 (2019).

26. Yang, S. et al. Crystal structure of the Frizzled 4 receptor in a ligand-free state. Nature 560, 666–670 (2018).

27. Nichols, A. S., Floyd, D. H., Bruinsma, S. P., Narzinski, K. & Baranski, T. J. Frizzled receptors signal through G proteins. Cell. Signal. 25, 1468–1475 (2013).

28. Liu, T. et al. G Protein Signaling from Activated Rat Frizzled-1 to the beta-Catenin–Lef-Tcf Pathway. Science 292, 1718–1722 (2001).

29. Koval, A. & Katanaev, V. L. Wnt3a stimulation elicits G-protein-coupled receptor properties of mammalian Frizzled proteins. Biochem. J. 433, 435–440 (2011).

30. Schulte, G. & Wright, S. C. Frizzleds as GPCRs – More Conventional Than We Thought! *Trends Pharmacol*. Sci. 1–15 (2018). doi:10.1016/j.tips.2018.07.001

31. Slusarski, D. C., Corces, V. G. & Moon, R. T. Interaction of Wnt and a Frizzled homologue triggers G-protein-linked phosphatidylinositol signalling. Nature 390, 410–413 (1997).

32. Panakova, D., Werdich, A. A. & Macrae, C. A. Wnt11 patterns a myocardial electrical gradient through regulation of the L-type Ca(2+) channel. Nature 466, 874–878 (2010).

33. López-López, J. R., Shacklock, P. S., Balke, C. W. & Wier, W. G. Local calcium transients triggered by single L-type calcium channel currents in cardiac cells. Science 268, 1042–1045 (1995).

34. Gomez-Ospina, N. et al. A Promoter in the Coding Region of the Calcium Channel Gene CACNA1C Generates the Transcription Factor CCAT. PLoS ONE 8, e60526 (2013).

35. Gomez-Ospina, N., Tsuruta, F., Barreto-Chang, O., Hu, L. & Dolmetsch, R. The C terminus of the L-type voltage-gated calcium channel Ca(V)1.2 encodes a transcription factor. Cell 127, 591–606 (2006).

36. Dijane, A., Riou, J.-F., Umbhauer, M., Boucaut, J.-C. & Shi, D.-L. Role of frizzled 7 in the regulation of convergent extension movements during gastrulation in Xenopus laevis. Development 127, 3091–3100 (2000).

37. Witzel, S., Zimyanin, V., Carreira-Barbosa, F., Tada, M. & Heisenberg, C.-P. Wnt11 controls cell contact persistence by local accumulation of Frizzled 7 at the plasma membrane. J. Cell Biol. 175, 791–802 (2006).

38. Katanaev, V. L., Ponzielli, R., Sémériva, M. & Tomlinson, A. Trimeric G protein-dependent frizzled signaling in Drosophila. Cell 120, 111–122 (2005).

39. Herbst, K. J., Allen, M. D. & Zhang, J. Spatiotemporally Regulated Protein Kinase A Activity Is a Critical Regulator of Growth Factor-Stimulated Extracellular Signal-Regulated Kinase Signaling in PC12 Cells. Mol. Cell. Biol. 31, 4063–4075 (2011).

40. Wong, W. & Scott, J. D. AKAP signalling complexes: focal points in space and time. Nat. Rev. Mol. Cell Biol. 5, 959–970 (2004).

41. Hundsrucker, C., Rosenthal, W. & Klussmann, E. Peptides for disruption of PKA anchoring. Biochem. Soc. Trans. 34, 472–473 (2006).

42. Huang, C.-J., Tu, C.-T., Hsiao, C.-D., Hsieh, F.-J. & Tsai, H.-J. Germ-line transmission of a myocardium-specific GFP transgene reveals critical regulatory elements in the cardiac myosin light chain 2 promoter of zebrafish. Dev. Dyn. 228, 30–40 (2003).

43. Dong, F., Feldmesser, M., Casadevall, A. & Rubin, C. S. Molecular characterization of a cDNA that encodes six isoforms of a novel murine A kinase anchor protein. J. Biol. Chem. 273, 6533–6541 (1998).

44. Merks, A. M. et al. Planar cell polarity signalling coordinates heart tube remodelling through tissue-scale polarisation of actomyosin activity. Nat Commun 9, 2161 (2018).

45. Schulte, G. European Journal of Pharmacology. European Journal of Pharmacology 1–5 (2015). doi:10.1016/j.ejphar.2015.05.031

46. Weivoda, M. M. et al. Wnt Signaling Inhibits Osteoclast Differentiation by Activating Canonical and Noncanonical cAMP/PKA Pathways. J Bone Miner Res 31, 65–75 (2015).

47. Ramaswamy, G. et al. Controls Cortical BoneQuality by Regulating OsteoclastDi erentiation via cAMP/PKA and. 1–11 (2017). doi:10.1038/srep45140

48. Matsuoka, A. et al. Progesterone Increases Manganese Superoxide Dismutase Expression via a cAMP-Dependent Signaling Mediated by Noncanonical Wnt5a Pathway in Human Endometrial Stromal Cells. The Journal of Clinical Endocrinology & Metabolism 95, E291–E299 (2010).

49. Manni, S., Mauban, J. H., Ward, C. W. & Bond, M. Phosphorylation of the cAMP-dependent protein kinase (PKA) regulatory subunit modulates PKA-AKAP interaction, substrate phosphorylation, and calcium signaling in cardiac cells. J. Biol. Chem. 283, 24145–24154 (2008).

50. Diviani, D., Dodge-Kafka, K. L., Li, J. & Kapiloff, M. S. A-kinase anchoring proteins: scaffolding proteins in the heart. Am. J. Physiol. Heart Circ. Physiol. 301, H1742–53 (2011).

51. Gao, T. et al. cAMP-dependent regulation of cardiac L-type Ca2+ channels requires membrane targeting of PKA and phosphorylation of channel subunits. Neuron 19, 185–196 (1997).

52. Dema, A. et al. The A-Kinase Anchoring Protein (AKAP) Glycogen Synthase Kinase 3 Interaction Protein (GSKIP) Regulates-Catenin through Its Interactions with Both Protein Kinase A (PKA) and GSK3 beta. J. Biol. Chem. 291, 19618–19630 (2016).

53. Lin, C. et al. Cypher/ZASP is a novel A-kinase anchoring protein. Journal of Biological Chemistry 288, 29403–29413 (2013).

54. Gold, M. G. et al. AKAP2 anchors PKA with aquaporin-0 to support ocular lens transparency. EMBO Mol Med 4, 15–26 (2012).

55. Thakkar, A., Aljameeli, A., Thomas, S. & Shah, G. V. A-kinase anchoring protein 2 is required for calcitonin-mediated invasion of cancer cells. Endocr. Relat. Cancer 23, 1–14 (2016).

56. Ulrich, F. et al. Slb/Wnt11 controls hypoblast cell migration and morphogenesis at the onset of zebrafish gastrulation. Development 130, 5375–5384 (2003).

57. Gordon, L. R., Gribble, K. D., Syrett, C. M. & Granato, M. Initiation of synapse formation by Wnt-induced MuSK endocytosis. Development 139, 1023–1033 (2012).

58. Lee, H., Lee, S. J., Kim, G.-H., Yeo, I. & Han, J.-K. PLD1 regulates Xenopus convergent extension movements by mediating Frizzled7 endocytosis for Wnt/PCP signal activation. Dev. Biol. 411, 38–49 (2016).

59. Gessert, S. & Kühl, M. The multiple phases and faces of wnt signaling during cardiac differentiation and development. Circulation research 107, 186–199 (2010).

60. Pandur, P., Läsche, M., Eisenberg, L. M. & Kühl, M. Wnt-11 activation of a non-canonical Wnt signalling pathway is required for cardiogenesis. Nature 418, 636–641 (2002).

61. Cohen, E. D., Tian, Y. & Morrisey, E. E. Wnt signaling: an essential regulator of cardiovascular differentiation, morphogenesis and progenitor self-renewal. Development 135, 789–798 (2008).

62. Mazzotta, S., et al. Stem Cell Reports. Stem Cell Reports 7, 764–776 (2016).

63. Mosimann, C. et al. Chamber identity programs drive early functional partitioning of the heart. Nat Commun 6, 8146 (2015).

64. Aye, T.-T. et al. Reorganized PKA-AKAP associations in the failing human heart. J. Mol. Cell. Cardiol. 52, 511–518 (2012).

65. Garcia-Gras, E. et al. Suppression of canonical Wnt/beta-catenin signaling by nuclear plakoglobin recapitulates phenotype of arrhythmogenic right ventricular cardiomyopathy. J. Clin. Invest. 116, 2012–2021 (2006).

66. Asimaki, A. et al. Identification of a new modulator of the intercalated disc in a zebrafish model of arrhythmogenic cardiomyopathy. Sci Transl Med 6, 240ra74 (2014).

67. Woo, L. A. et al. High-content phenotypic assay for proliferation of human iPSC-derived cardiomyocytes identifies L-type calcium channels as targets. J. Mol. Cell. Cardiol. 127, 204–214 (2018).

68. Weinberger, F., Mannhardt, I. & Eschenhagen, T. Engineering Cardiac Muscle Tissue: A Maturating Field of Research. Circulation research 120, 1487–1500 (2017).

69. Fujita, B. & Zimmermann, W.-H. Myocardial tissue engineering strategies for heart repair: current state of the art. Interact Cardiovasc Thorac Surg 27, 916–920 (2018).

70. Ronaldson-Bouchard, K. et al. Advanced maturation of human cardiac tissue grown from pluripotent stem cells. Nature 556, 239–243 (2018).

71. Heisenberg, C. P. et al. Silberblick/Wnt11 mediates convergent extension movements during zebrafish gastrulation. Nature 405, 76–81 (2000).

72. Schwanhäusser, B. et al. Global quantification of mammalian gene expression control. Nature 473, 337–342 (2011).

73. Livak, K. J. & Schmittgen, T. D. Analysis of relative gene expression data using real-time quantitative PCR and the 2(-Delta Delta C(T)) Method. Methods 25, 402–408 (2001).

74. Rappsilber, J., Mann, M. & Ishihama, Y. Protocol for micro-purification, enrichment, pre-fractionation and storage of peptides for proteomics using StageTips. Nat Protoc 2, 1896–1906 (2007).

75. Burger, A. et al. Maximizing mutagenesis with solubilized CRISPR-Cas9 ribonucleoprotein complexes. Development 143, 2025–2037 (2016).

76. Lindsay, H. et al. CrispRVariants charts the mutation spectrum of genome engineering experiments. Nat. Biotechnol. 34, 701–702 (2016).

77. Thisse, C. et al. High-resolution in situ hybridization to whole-mount zebrafish embryos. Nat Protoc 3, 59–69 (2008).

78. Chernyavskaya, Y. et al. Voltage-gated calcium channel CACNB2 (β2.1) protein is required in the heart for control of cell proliferation and heart tube integrity. Dev. Dyn. 241, 648–662 (2012).

79. Hohendanner, F. et al. Intracellular dyssynchrony of diastolic cytosolic [Ca²⁺] decay in ventricular cardiomyocytes in cardiac remodeling and human heart failure. Circulation research 113, 527–538 (2013).

80. Heinzel, F. R. et al. Spatial and temporal inhomogeneities during Ca2+ release from the sarcoplasmic reticulum in pig ventricular myocytes. Circulation research 91, 1023–1030 (2002).

81. Cox, J. & Mann, M. MaxQuant enables high peptide identification rates, individualized p.p.b.-range mass accuracies and proteome-wide protein quantification. Nat. Biotechnol. 26, 1367–1372 (2008).

